# Structural principles of B-cell antigen receptor assembly

**DOI:** 10.1101/2022.08.13.503858

**Authors:** Ying Dong, Xiong Pi, Frauke Bartels-Burgahn, Deniz Saltukoglu, Zhuoyi Liang, Jianying Yang, Yumei Zheng, Frederick W. Alt, Michael Reth, Hao Wu

**Affiliations:** Department of Biological Chemistry and Molecular Pharmacology, Harvard Medical School, Boston, MA, USA; Program in Cellular and Molecular Medicine, Boston Children’s Hospital, Boston, MA, USA; Institute of Biology III, Faculty of Biology, University of Freiburg, Freiburg, Germany; Signaling Research Centers CIBSS and BIOSS, University of Freiburg, Freiburg, Germany; HHMI, Boston Children’s Hospital, Boston, MA, USA; Department of Genetics, Harvard Medical School, Boston, MA, USA

## Abstract

The B-cell antigen receptor (BCR) is composed of a membrane-bound immunoglobulin (mIg) of class M, D, G, A or E for antigen recognition and a disulfide-linked heterodimer between Igα and Igβ (Igα/β, also known as CD79A and CD79B) that functions as the signalling entity. The organizing principle of BCR assembly remains elusive. Here we report the cryo-electron microscopy structures of the intact IgM class BCR at 8.2 Å resolution and its Fab-deleted form (IgM BCRΔFab) at 3.6 Å resolution. At the ectodomain (ECD), Igα and Igβ position their respective Ig folds roughly in parallel with an approximate 2-fold symmetry, which is distinct from structures of Igβ/β homodimers. Unlike previous predictions, the BCR structure displays an asymmetric arrangement, in which the Igα/β ECD heterodimer mainly uses Igα to associate with Cµ3-Cµ4 domains of one heavy chain (µHC) while leaving the other heavy chain (µHC’) empty. The transmembrane domain (TMD) helices of the two µHCs also deviate from the 2-fold symmetry of the Cµ3-Cµ4 domain dimer and form together with the TMD helices of the Igα/β heterodimer a tight 4-helix bundle. The asymmetry at the TMD helices prevents the recruitment of two Igα/β heterodimers. Surprisingly, the connecting peptides (CPs) between the ECD and TMD are braided together through striking charge complementarity, resulting in intervening of the CP of µHC in between those of Igα and Igβ and crossover of the TMD relative to ECD for the Igα/β heterodimer, to guide the TMD assembly. Interfacial analyses suggest that the IgM BCR structure we present here may represent a general organizational architecture of all BCR classes. Our studies thus provide a structural platform for understanding B-cell signalling and for designing rational therapies against BCR-mediated diseases.

---

B cells can recognize structurally diverse antigens. Upon antigen recognition, B cells can be activated and differentiated into plasma cells that secrete antibodies to neutralize the antigens. Antigen recognition by B cells is fulfilled by mIg in BCR, as well as secreted Ig (sIg, or antibody), differing only at the C-terminal end by alternative mRNA processing^1–3^. The different classes or isotypes of mIg and sIg − IgM (µ), IgD (**δ**) IgG (γ), IgA (α) or IgE (ε) – have different constant regions. IgM is the first isotype expressed on all immature and naïve mature B cells. Like the basic architecture of an antibody, mIg comprises a symmetric homodimer of a heterodimer of membrane-bound heavy chain and light chain. The BCR signalling components Igα and Igβ each carry an immunoreceptor tyrosine-based activation motifs (ITAM) at their cytoplasmic tails. Despite the importance of the BCR in B-cell development and activation, no structural information of an intact BCR has been reported.

## Reconstitution and structure determination

To pursue structure determination of a BCR by cryo-electron microscopy (cryo-EM), we utilized a derivative of the murine myeloma cell line J558L (J558Lµm15-25/mb-1NFlag) that stably co-expresses the mIgM heavy and lambda light chains together with a FLAG-tagged Igα-YFP fusion protein and wild-type Igβ (Fig. 1a, Extended Data Fig. 1). The intact IgM BCR was purified by anti-FLAG affinity and gel filtration chromatography, followed by SDS-PAGE showing the co-expression of all the individual components (Extended Data Fig. 2). Two-dimensional (2D) classification (Extended Data Fig. 3) of the cryo-EM data showed large-scale conformational changes at the Fab region due to the known flexibility at the mIgM hinge region^4–7^. The cryo-EM analysis produced a density map at 8.2 Å resolution (Fig. 1b and Extended Data Fig. 4).

**Fig. 1.**
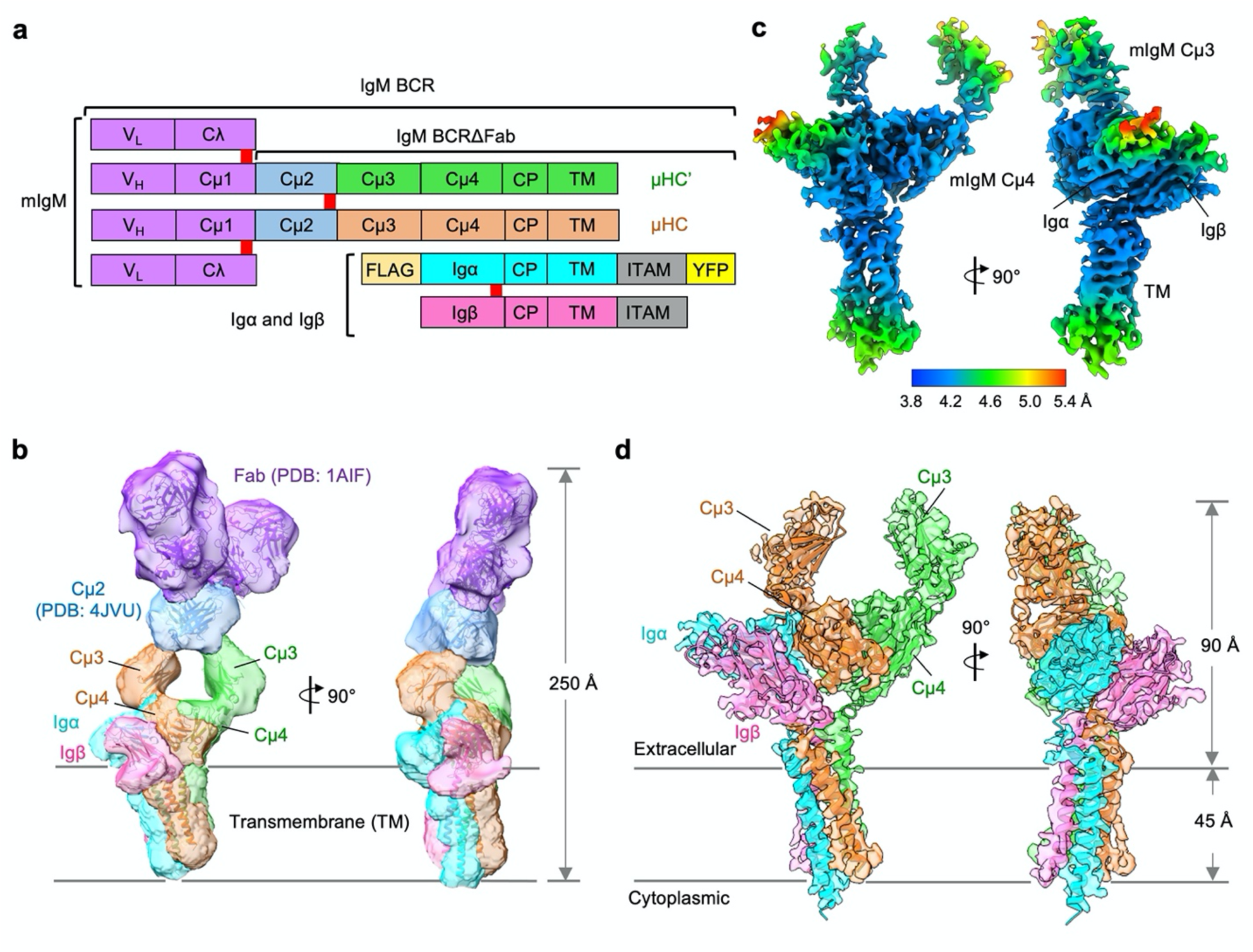
Cryo-EM maps of the IgM-BCR. **a**, Domain organization of the IgM BCR shown with Fab in purple, Cµ2 in blue, Cµ3-Cµ4-CP-TM in green and orange, Igα-CP-TM and Igβ-CP-TM in cyan and pink, followed by the ITAM-containing cytoplasmic tail shown in gray. Disulfide bonds are shown as red rectangles. FLAG tag and YFP fusions to Igα are shown in light orange and yellow, respectively. mIgM heavy chains µHC and µHC’ are proximal and distal to the Igα/β heterodimer, respectively. CP: connecting peptide. **b**, Cryo-EM map of full-length IgM BCR at 8.2 Å resolution (contour level: 0.5 σ) superimposed with the model. Fab (PDB: 1AIF) and mIgM-Cµ2 (PDB:4JVU) fitting into the cryo-EM density map. **c**, Local resolution distribution of the IgM BCRΔFab map at 3.6 Å resolution (contour level, 3.0 σ). **d**, Cryo-EM map of IgM BCRΔFab (contour level, 3.0 σ) superimposed with the model.

To optimize the overall resolution of the BCR structure, we modified the cell line to express a truncated mIgM lacking the two Fab arms, and purified BCRΔFab using the same procedure used for intact BCR (Extended Data Fig. 1, 2). 2D classification of the cryo-EM data revealed clearer structural features at both the ECD and the TMD in comparison with intact BCR (Extended Data Fig. 3). After 3D classification and focused refinement, a homogeneous dataset of ∼400,000 particles produced a final map at an overall resolution of 3.6 Å with best densities at Cµ4, the TMD bundle and part of Igα and Igβ (Fig. 1c, Extended Data Fig. 5a, Table 1). The particles angular distribution and Fourier shell correlation (FSC) plots are shown (Extended Data Fig. 5b, c). We used AlphaFold running on ColabFold notebook^8,9^ to assist model generation and density fitting (Extended Data Fig. 6, 7). Because the mIgM Cµ2 domains and the cytosolic domain of the Igα/β heterodimer are not visible due to their previously recognized dynamic nature^10^, the final model of BCRΔFab includes regions of mIgM from Cµ3 to TMD and regions of the Igα/β heterodimer up to the TMD (Fig. 1d). Fitting of the BCRΔFab structure together with the crystal structures of Fab and Cµ2 domains^11,12^ into the density led to the model of full-length IgM BCR (Fig. 1b). The linkers between the ECD and the TMD in mIgM, Igα and Igβ turned out to be important for the organization of the BCR (see below), and we named them connecting peptides (CPs), following the domain nomenclatures used to describe the T-cell antigen receptor (TCR)^10^.

## The ECD interaction in the Igα/β heterodimer

Igα and Igβ ECDs both assume an Ig fold, and interact with each other with their β-sandwich domains roughly in parallel (Fig. 2a-c). Despite the low sequence identity (22%), Igα and Igβ are highly similar in structures with RMSD of 3.2 Å (Fig. 2d). They are related by an approximate 2-fold axis (166 ° in rotation) and linked by a disulfide bond at the interface between two equivalent residues, Igα C113 and Igβ C135 (Fig. 2b, c), which is consistent with a previous prediction^13^. There are 5 glycosylation sites on Igα and Igβ, all of which had cryo-EM densities for the first N-acetylglucosamine (NAG) residue in their N-linked glycans (Fig. 2b). The glycan on N68 of Igα situates at the interface among Igα, Igβ and mIg, suggesting that it may play a role in the assembly of the BCR.

**Fig. 2.**
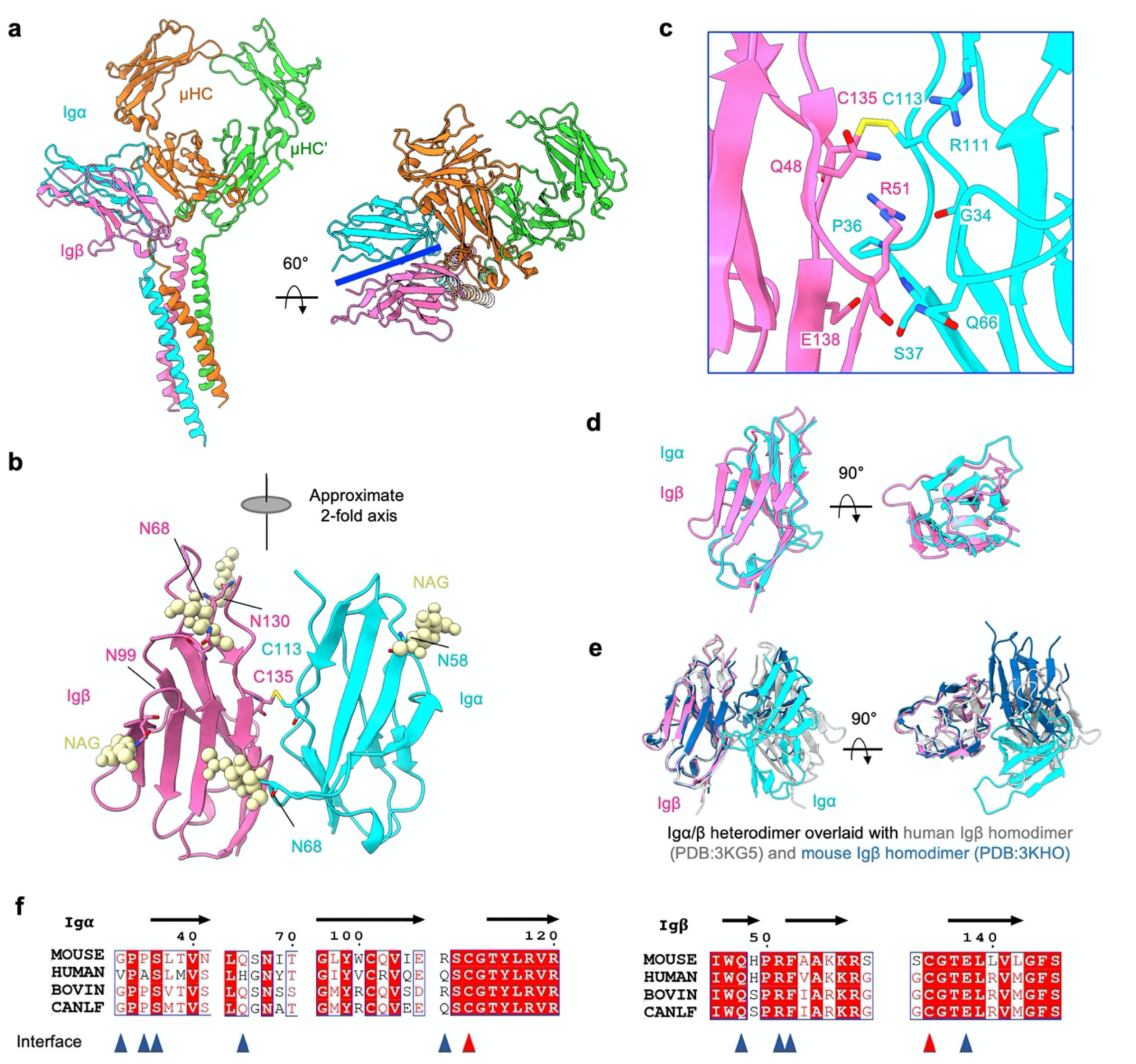
Interactions between the Ig domains of Igα and Igβ.Kardon. **a**, The model of IgM BCRΔFab shown in ribbon with different views. The ECD interface between Igα and Igβ is represented by a blue line. **b**, Ribbon diagram of the Ig domains of the Igα/β heterodimer. The intersubunit disulfide bond between Igα C113 and Igβ C135 and the observed N-linked glycans are shown. The Ig domains of Igα and Igβ are related by an approximate 2-fold axis. **c**, Detailed interfacial interactions. **d**, Alignment between the Ig domains of Igα and Igβ, showing structural similarity. **e**, Alignment of Ig domains of the Igα/β heterodimer with two different crystal structures of the Igβ/β homodimer, showing marked differences. **f**, Alignment of part of the Igα and Igβ ECD protein sequences from different species. Interacting amino acids within the Igα/β heterodimer are marked by triangle symbols, Igα C113 and Igβ C135 are marked in red. CANLF: Dog.

Igα and Igβ interact extensively at the ECD with a buried surface area of ∼500 Å^2^ each as calculated on the PDBePISA server^14^. The interaction is rich in potential hydrogen bonds, such as those among Igβ E138 and the amides of Igα S37 and Igβ P36, between Igβ R51 and the carbonyl oxygen atoms of Igα G34, between Igα R111 and Igβ Q48, and between Igα Q66 and carbonyl oxygen atom of Igβ R51 (Fig. 2c). Of note, the packing of the Igα/β heterodimer is distinct from structures of Igβ/β homodimers which are also linked by the analogous disulfide bond at Igβ C135 ^13^ (Fig. 2e). Residues at the interface between Igα and Igβ are largely conserved across species (Fig. 2f), suggesting an evolutionarily conserved interaction.

## The ECD interaction of the Igα/β heterodimer with mIgM

The Igα/β ECD interacts with the ring-shaped and symmetric Cµ3-Cµ4 dimer of mIgM mainly at one side, creating an asymmetry in the BCR with µHC proximal and µHC’ distal to Igα/β (Fig. 3a). In this interaction, Igα contributes ∼830 Å^2^ buried surface area and Igβ only contributes ∼90 Å^2^. Most of the contacts are with the Cµ4 domain of µHC with ∼750 Å^2^ buried surface area, in contrast to 170 Å^2^ at Cµ3 (Fig. 3a). The symmetric Cµ3-Cµ4 homodimer aligns well with the cryo-EM structure of sIgM at this region^15,16^ (Extended Data Fig. 8). The mutual interaction between the Igα/β heterodimer and mIgM has an electrostatic component shown by the largely negatively charged surface of the Cµ4 dimer and the largely positively charged surface of the Igα/β Complex (Fig. 3b). In particular, Cµ4 E428 forms an ionic pair with Igα R96 and a potential hydrogen bond with the amide of Igα Q66, and Cµ4 D432 and T412 are within salt bridge and hydrogen bonding interactions, respectively, to Igβ K55 (Fig. 3c). Potential hydrogen bonds are also present between Cµ4 H424 and Igα Y99, between Cµ4 R429 or Cµ4 P331 carbonyl oxygen atom and Igα N94, between Cµ4 Q366 and Igα T70, and between Cµ4 G371 carbonyl oxygen atom and Igα S67 (Fig. 3c). The glycan on Igα N68 and Igβ N99 may further enhance the Igα−Cµ4 interaction (Fig. 3c). In line with this is the different glycosylation of Igα when incorporated within the IgM BCR complex^17,18^. In addition to µHC that provides most of the interactions, Cµ4 E341 from µHC’ appears to form an ionic pair with Igα K93 (Fig. 3c). When the TCRβ constant Ig domain of the TCR^10^ is aligned with the Cµ4 domain of µHC, the CD3ε/γ signalling component of the TCR is situated at the same region as the Igα/β heterodimer (Fig. 3d).

**Fig. 3.**
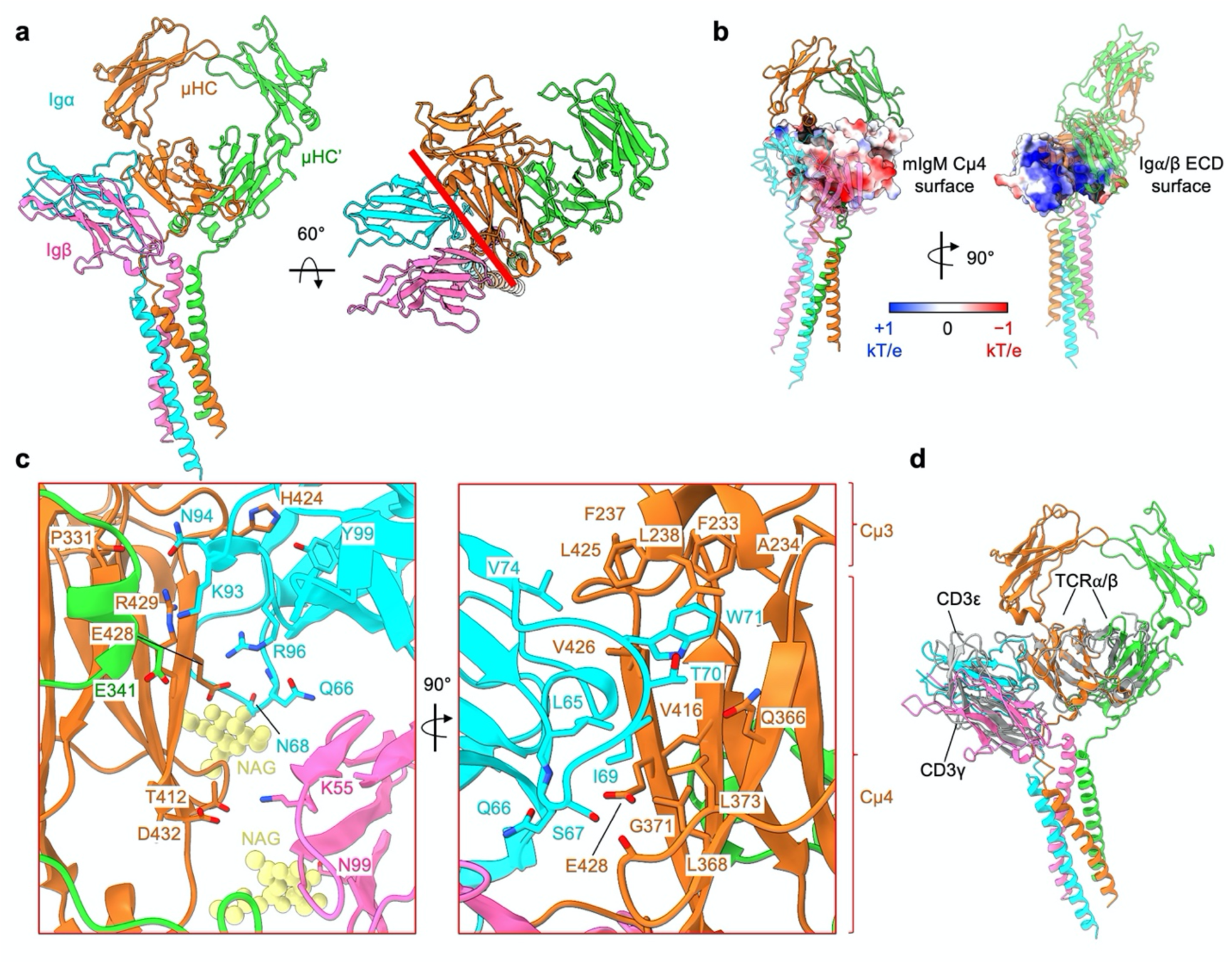
Ig domain interactions between the Igα/β heterodimer and mIgM. **a**, Model of the IgM BCRΔFab shown as ribbon diagram with different views. The Ig domain interface between Igα/β and mIgM is indicated with a red line. **b**, Electrostatic surfaces (−1 to +1 kT/e) of the interacting Cµ4 (left) and the Ig-domains of the Igα/β heterodimer (right). **c**, Detailed view of the interactions at the interface. Shown in yellow are the N-linked glycans on Igα (N99) and Igβ (N68) which may enhance the interaction. **d**, Alignment of the Cµ4 domain of the IgM BCR with the TCRβ constant domain (grey), showing that the Ig domains of the Igα/β and CD3ε/γ (grey, PDB: 6JXR) heterodimers occupy the similar location relative to mIgM and TCRα/β, respectively.

The sequence alignment of the ECD of different mIg isotypes shows that the key residues involved in the interactions with the Igα/β heterodimer are poorly conserved (Extended Data Fig. 9). In line with this observation is the finding that the isolated ECD of the Igα/β heterodimer binds strongly to a soluble Cµ3-Cµ4 construct of IgM but barely to that of the other isoforms^13^. However, as all mIg isotypes interact with the Igα/β heterodimer, the CP and/or TMD parts may play a dominant role for the stability of the BCR (see below).

## The CP interaction between mIgM and the Igα/β heterodimer

The BCR structure shows that the CPs of µHC, µHC’, Igα and Igβ intertwine as if being braided together as they connect to the TMD, and may play important specificity and energetic roles in BCR assembly (Fig. 4a). First of all, there is an apparent crossover by the CPs of Igα and Igβ so that their ECDs and TMDs switch positions (Fig. 4a). Interestingly, similar ECD to TMD crossovers are features of all TCR components, including the CD3**ε**/**γ**, CD3**ε**’/**δ** and TCRα/β complexes (Fig. 4a). However, unlike in TCRα/β, the ECDs and TMDs of µHC and µHC’ do not switch positions. The CP of µHC plays a dominant role in Igα/β binding and intervenes in between the CPs of Igα and Igβ, while the CP of µHC’ interacts only at the outer sides of the CP of Igβ and Cµ4 domain of µHC (Fig. 4a).

**Fig. 4.**
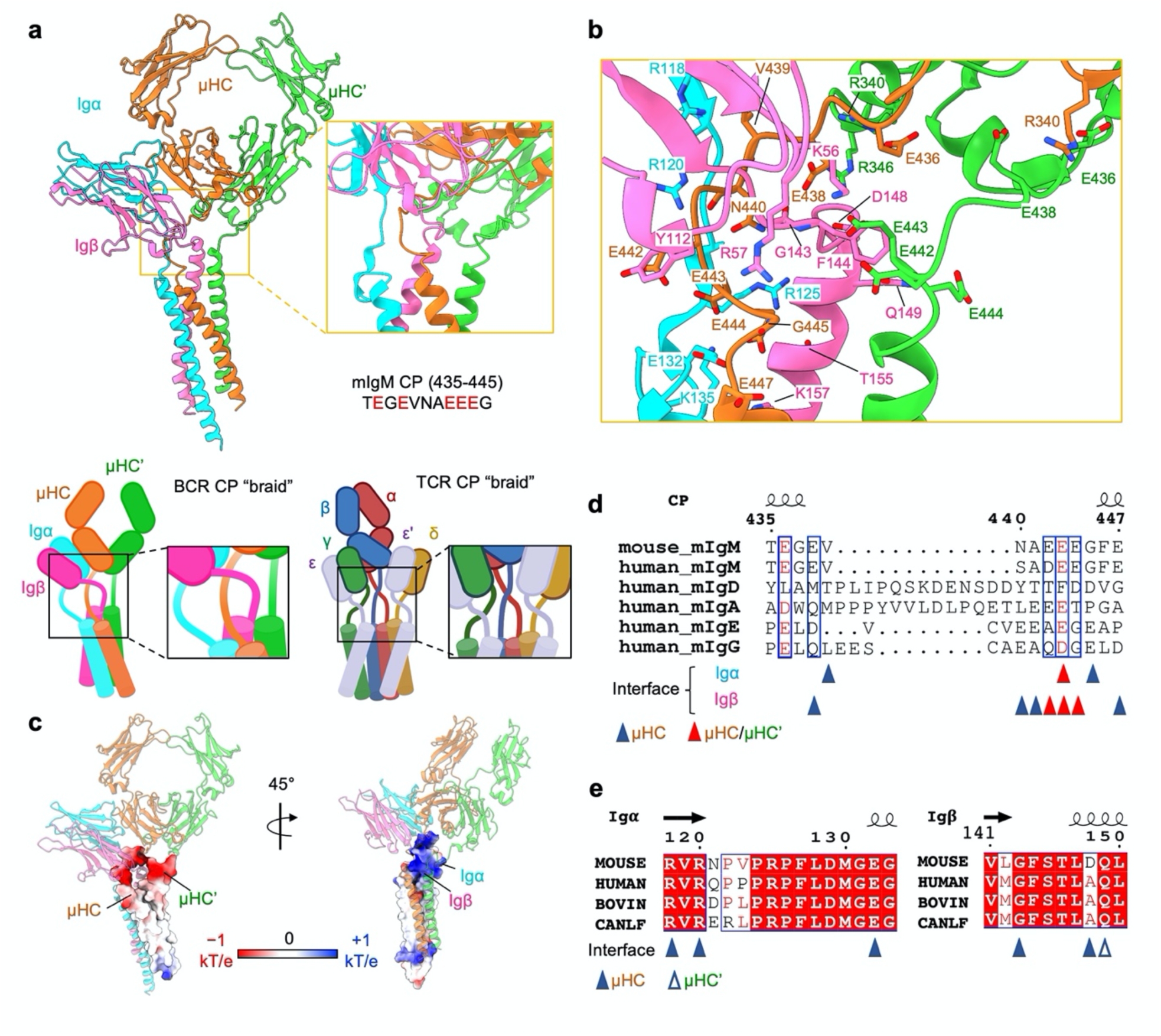
IgM BCR assembly at the CP regions. **a**, Global view (left) and enlarged view (right) of the CP region of the IgM BCR, showing the intertwining in this region. The sequence of murine mIgM CP (435-445) is shown with acidic residues in red. The assembly of BCR and TCR CPs is shown in schematics. **b**, Detailed interactions between the charged CP residues of the Igα/β heterodimer and the mIgM molecule. **c**, Electrostatic surfaces (−1 to +1 kT/e) at the CPs of mIgM (left) and the CPs of Igα and Igβ (right). **d**, CP sequences of different mIg isotypes. Acidic residues are boxed in blue rectangles. Residues in the interface of Igα/β with µHC and µHC’ are indicated by triangles. **e**, Sequence alignment of Igα/β CPs. Residues in the interface of Igα/β with µHC and µHC’ are indicated by triangles.

The mIgM CP contains a string of acidic residues (Fig. 4a, b) and displays strong negative electrostatic potential (Fig. 4c). By contrast, the surrounding region from Igα and Igβ including their CPs is highly positively charged (Fig. 4c). For the CP of µHC, specific interactions may include E436 with µHC’ R340, E438 with amide of Igβ G143 and F144, E442 with Igβ Y112, E443 with Igα K135, E444 with Igβ K157, and E447 with Igβ R57 (Fig. 4b). For the CP of µHC’, specific interactions may include E436 and E438 with µHC R340, E442 and E443 with Igβ K56, and E444 with Igβ Q149 (Fig. 4b). In addition, Igβ CP residue D148 interacts with µHC’ R346; for Igα, specific interactions may include R118 and R120 with the carbonyl oxygen atoms of µHC V439 and N440 respectively, R125 with Igβ T155, and E132 with the amide of µHC G445 (Fig. 4b).

Collectively, these data suggest that the CPs braid together through largely electrostatic interactions, and that they may be crucial for guiding the assembly of the 4-helix bundle in the TMD. Sequence alignment among the CP regions of Igα, Igβ, and different isotypes of mIg showed that the interacting residues are largely conserved (Fig. 4d, e), which further support a role of the CP for BCR assembly. However, the CPs of mIg isotypes differ in their lengths, suggesting potential variations in their lateral interactions. For example the CP of mIgD contains 8 additional residues and this may be related to the higher stability of the IgD BCR complex in comparison the IgM BCR^19^.

## The TMD 4-helix bundle assembly

The four TMD helices assemble into a tight 4-helix bundle (Fig. 5a), burying an average of 1300 Å^2^ surface area per helix, and thus a massive total surface area of ∼5200 Å. The Igα and Igβ subunits differ in their contact with TMDs of mIgM. Igα contacts with TMD helices of both µHC and µHC’, whereas Igβ only contacts with that of µHC’ (Fig. 5b). Overall, the TMD residues in close proximity to the outer or inner surface of the plasma membrane are often polar or charged while the core regions of the helices contain mainly hydrophobic residues (Fig. 5b-g), consistent with the generic organization of most TMD helices. Outer membrane-proximal charged or polar interactions include those among Igβ Q149, R152 and R153 and µHC’ E447, among Igβ K157, Igα N136 and R137, and µHC N448 and T452, between Igα K135 and Igβ N154 (Fig. 5b, top left). At the cytoplasmic side, membrane-proximal charged or polar interactions include those among Igα R160 and K161, and Igβ D181, K182 and D183 (Fig. 5b, lower right).

**Fig. 5.**
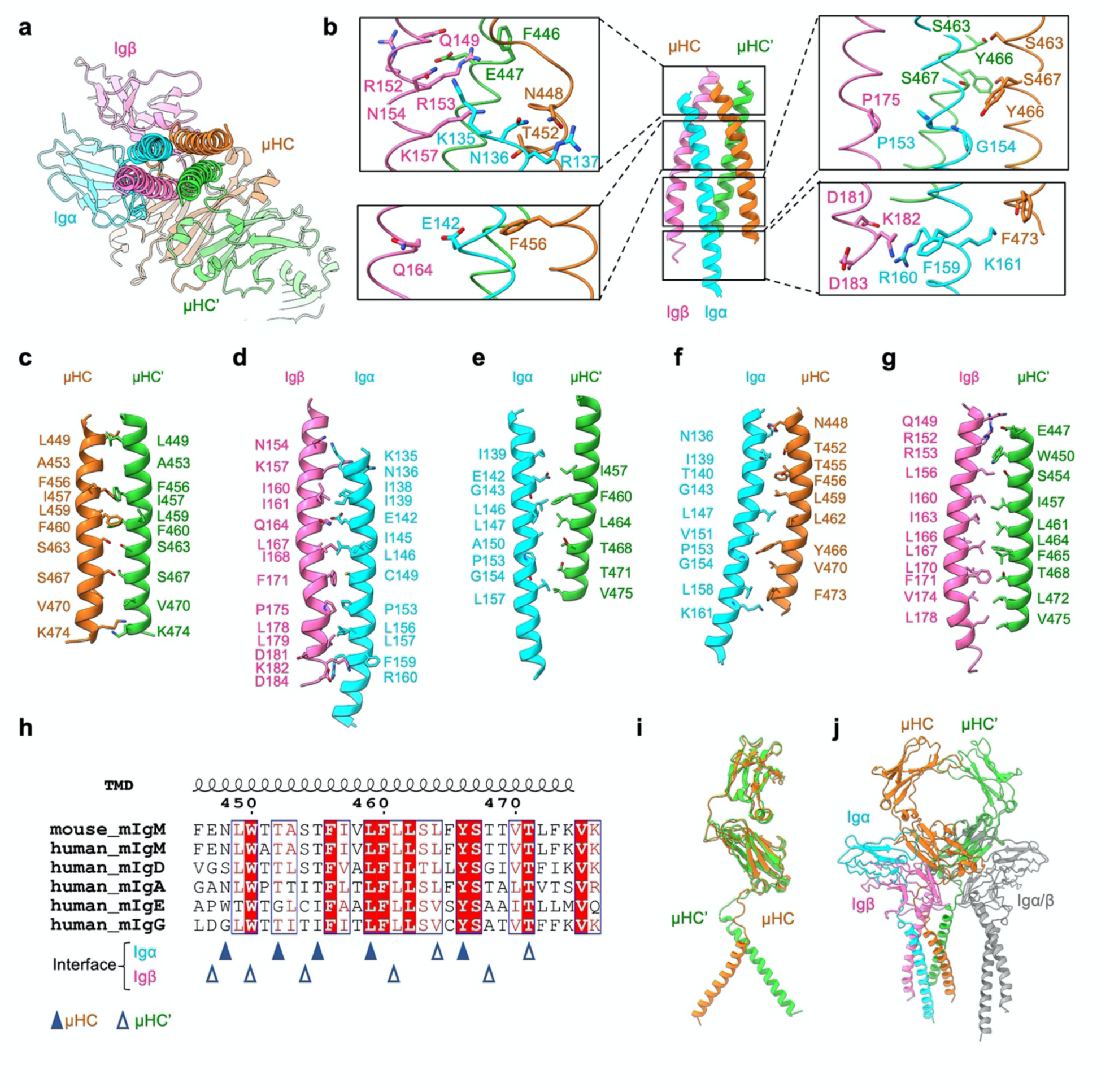
TMD assembly of the IgM BCR. **a**, Bottom view of the TMD assembly below the ECD of IgM BCR shown in ribbon. **b**, Key polar interactions at the TMD including those at the E/Q-X_10_-P motif on Igα and Igβ and YS motif on mIgM. **c-g**, Interactions between pairs of helices of TMD. **h**, Residues in mIgM mediating TMD helical bundle formation are partly conserved among different mIg isotypes. **i**, Structural alignment of the µHC and µHC’ in an mIgM dimer, showing the different, asymmetric conformation at the TMD. **j**, Structurally aligning a second Igα/β heterodimer to the empty side of the mIgM Cµ4 dimer showed lack of interaction with the 4-helix bundle of TMD, which may explain why the second Igα/β heterodimer is not recruited to BCR.

However, the cryo-EM structure reveals that there are also charged or polar residues within the core region of the TMD. In particular, it is intriguing that Igα E142 is largely buried within the 4-helix bundle and surrounded by hydrophobic residues (Fig. 5d). One possibility is that E142 may form an anion-aromatic interaction with the absolutely conserved mIg residue F456 (Fig. 5b, lower left and 5h) as aromatic ring edge is known to exhibit positive electrostatic potential^20,21^. Another possibility is that the hydrophobic environment potentiates protonation of the E142 carboxylate. In addition, Igα E142 is adjacent to the equivalent residue Igβ Q164, which may further help to dissipate the charge. We hypothesize that these polar interactions within the membrane may help to precisely assemble the TMD, explaining the conservation of both Igα E142 and mIg F456 (Fig. 5h, Extended Data Fig. 10). The conservation of the charged Igα Residue within the membrane is reminiscent of the TCR helical bundle in which charged interactions help to bring the TMD helices together at the center of the membrane^10^.

Another hydrophilic cluster within the membrane involves Y466 and S467 of mIgM (Fig. 5b, top right). Here, a potential hydrogen bond is formed between two S467 residues of µHC and µHC’. The Y466 residue of µHC and that of µHC’ are distant from each other and only µHC Y466 forms a potential hydrogen bond with main chain of Igα G154, and packs against Igα P153, but not the equivalent Igβ P175 residue. In comparison with the poor conservation of the ECD interaction between the Igα/β heterodimer and different isoforms of mIg (Extended Data Fig. 9), the TMD domain interaction is largely conserved (Fig. 5h). The TMD helices of µHC and µHC’ are assembled asymmetrically as shown by their different positioning when superposed by the ECD (Fig. 5i). Consistently, if we add another Igα/Igβ heterodimer to the empty side of the mIg dimer, the TMD of the heterodimer does not contact the 4-helix bundle (Fig. 5j). This lack of interaction at the TMD may explain the 1:1 stoichiometry of the monovalent IgM BCR complex.

## Discussion

Our high-resolution cryo-EM structure of the murine IgM BCR reveals a 1:1 mIgM-Igα/β complex that confirms previous biochemical and FRET data^22,23^. The mIgM molecule is a symmetric homodimer and it was previously thought that the IgM BCR complex also has a symmetric structure but this is clearly not the case. Most interactions for the stability of the complex are mediated by only one of the two mIgM heavy chains, namely µHC, whereas Igα plays a dominant role in the binding of the Igα/β heterodimer to the mIgM molecule. The latter finding explains why only the Igα/β heterodimer but not the previously described Igβ/β homodimer becomes part of the IgM BCR complex. A closer inspection of the receptor complex reveals that the symmetry is broken by the CP and TMD region of the mIgM molecule.

A comparison of our IgM BCR cryo-EM structure with that of the TCRα/β−CD3 complex reveals common and diverging features of these two important receptors of the adaptive immune system. Both receptors display an asymmetric assembly with the signalling subunits tagged to one side of the antigen binding component. The intertwined CP regions and the close interactions of the α-helical TMDs of the subcomponents play an important role for the stability of both receptors. For TCR, charged interactions deep within the TMD helices are critical for its assembly^10^. For BCR, the evolutionary conserved E/Q-X_10_-P motifs in Igα (E142-X_10_-P153) and Igβ (Q164-X10-P175) and the YS motifs (Y466-S467) in mIg define certain polar interactions within the membrane, albeit also asymmetrically instead of symmetrically as predicted^24,25,26,27^. The TCRα/µ incorporates three CD3 signalling dimers and thus forms an 8-helix bundle in the TCR TMD, with the α/β helices surrounded by helices of the CD3 components. By contrast, the IgM BCR forms a 4-helix bundle TMD with the µHC and µHC’ helices exposed on one side to the lipid bilayer.

A previous analysis of mIg TMD sequences suggested that the side of the α-helix highly conserved in all mIg isotypes (TM-C) associates with TMDs of the Igα/β heterodimer, whereas the other side harbouring class-specific residues (TM-S) is involved in µHC dimerization and mIg oligomerization^24^. Our cryo-EM structure shows that this assignment is not correct as the central TM-C side mediates µHC dimerization and its neighbouring residues dominantly interact with Igα TMD and to a lesser extent Igβ TMD. What then is the role of the class-specific residues at the TM-S side? In the case of the IgD BCR, these residues were shown to involve in BCR oligomerization and that their mutation results in receptor activation, supporting the dissociation activation model (DAM) of BCR activation^28^ (Fig. 6, Extended Data Fig. 11). However, the conserved residues at the TM-S side may also be involved in the binding to lateral interactors of the BCR. One feasible candidate is the transmembrane phosphatase CD45 that controls the resting state of T and B cells^29^ (Fig. 6, Extended Data Fig. 11). Another possible interactor is CD19 that is found in close proximity to the IgM BCR only on activated B cells^29^ (Fig. 6, Extended Data Fig. 11), and antigen binding and dissociation of the IgM BCR oligomer could expose one side of the mIgM for binding to CD19.

**Fig. 6.**
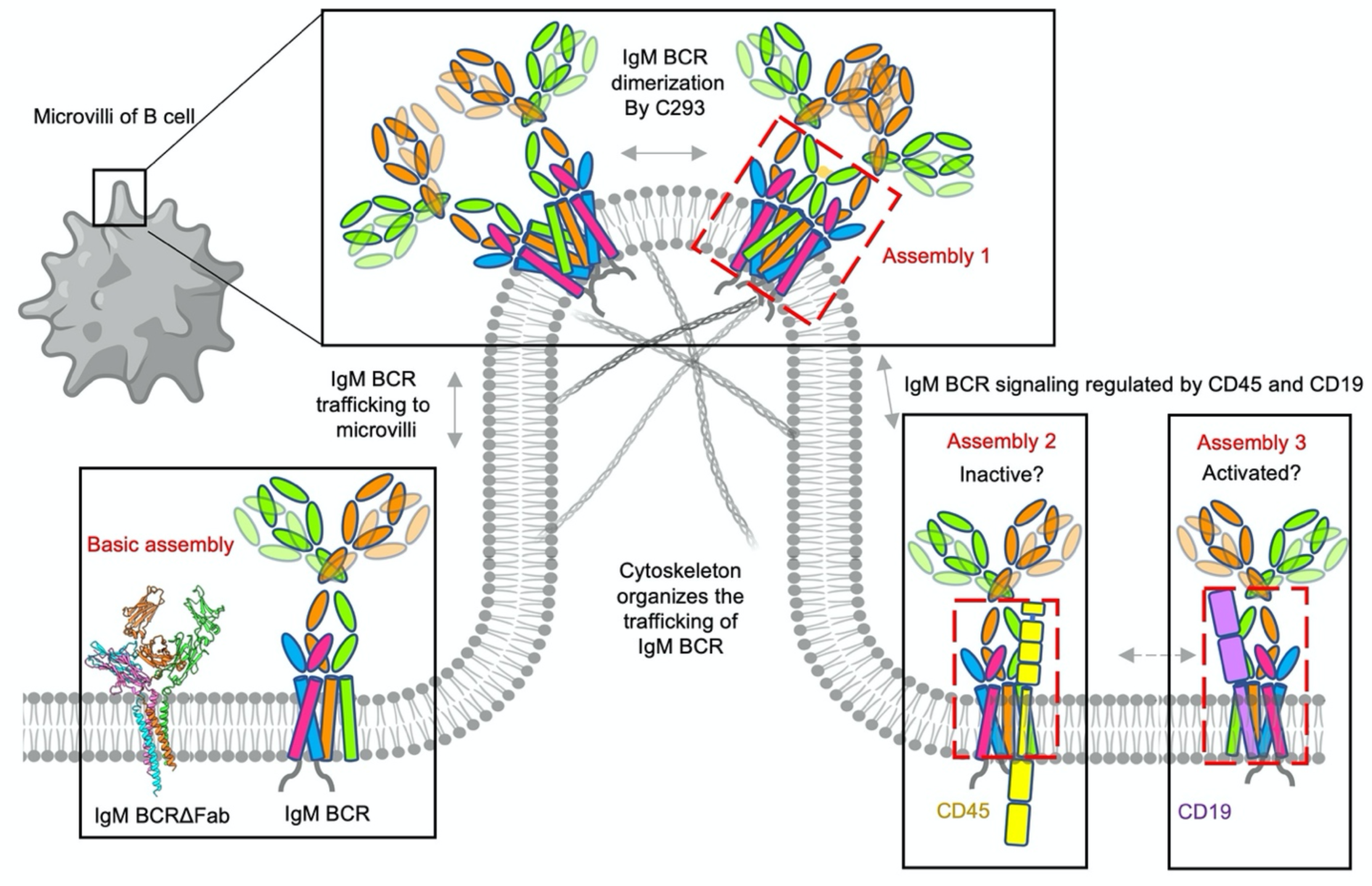
Proposed models for dynamic assembly of IgM BCR with itself and other signalling components. The basic assembly unit is the IgM BCR observed in this study. Once trafficked to the microvilli of B cells, IgM BCR may dimerize into a homodimer. In addition, IgM BCR may associate with CD45 or CD19 under different conditions.

Unfortunately, none of the now available cryo-EM structures of TCR and BCR revealed the conformation of the cytoplasmic tails of the signalling subunits. It thus remains unclear how antigen binding is transduced to the cytoplasmic ITAM tyrosine residues for accessibility and phosphorylation. We hypothesize that the surprising CP crossover in BCR assembly, which is also a feature for TCR, may allow a scissor-like process that coverts a slight movement of the ECD after antigen binding to a major impact on the conformation of the TMD and cytoplasmic receptor tails with a low energetic cost. The cytoplasmic tails of the CD3 subunits are positively charged, which may associate with the negatively charged inner leaflet of the plasma membrane in the resting state to prevent their phosphorylation^30^. However, the cytoplasmic tails of Igα/β are dominantly negatively charged, whose membrane binding may require positively charged intermediate proteins. This could, for example, be the highly positively charged BAR domain proteins that are associated with curved membranes^31,32^. A recent study showed that oligomeric IgM BCR clusters on living B cells are associated with a network of ridges and microvilli that both contain curved membranes^33^ (Fig. 6). How the cytoplasmic tails of the TCR and the BCR are organized in the resting state and altered during the activation process is clearly an important issue of further structural studies.

## Materials and methods

### Construct design

The Flag-Igα-YFP construct encodes a chimeric Igα protein with the FLAG-tag sequence inserted after the leader peptide of Igα. The cytoplasmic tail of the chimeric Igα is truncated 10 aa after the TMD at G169 and fused with YFP through a RSIATRS linker. The cDNA of the Flag-Igα-YFP construct is cloned into the retroviral vector pMOWS. The expression vector for the truncated IgM BCRΔFab (pMS-RL2b) was derived from a pMOWS expressing murine mIgM heavy chain by replacing the sequence encoding the VH and Cµ1 domains with a Strep tag (WSHPQFEK) connected to the Cµ2 domain via a rigid linker of the sequence A(EAAAK)3A.

### Generation of stable cell lines

The cell line J558Lµm15-25 expresses a murine mIgM molecule with specificity for the hapten 4-hydroxy-5-iodo-3-nitrophenyl acetyl (NIP), the WT murine Igβ but no Igα. For the expression of a fully assembled IgM BCR complex on the cell surface the J558Lµm15-25 cells were transduced with a retroviral vector encoding the Flag-Igα-YFP construct. The transfectants were sorted for YFP positive cells and the expression of a NIP-specific IgM BCR on the cell surface using an APC-coupled 1-NIP peptide and collected by flow cytometry. For the generation of the IgM BCRΔFab expressing cell, we transfected the pMS-RL2b vector into the J558L/Flag-Igα-YFP clone 8A1. Before large scale expression and purification, stable cell lines were further sorted by flow cytometry to collect cells with the highest YFP expression.

### Flow Cytometry

Flow cytometric analysis was used for measuring the co-expression of mIgM and Igα/β in IgM BCR and IgM BCRΔFab cells. Cells were collected and washed once with PBS. Then, the cells were stained for the surface makers with or without the indicated antibodies (IgM-APC). The APC-IgM antibody was diluted 100 times from stock concentration at room temperature for 5 mins. The stained cells were washed once with PBS and resuspended in PBS before flow cytometric analysis. Attune Flow Cytometers (Thermo) was used for flow cytometry. Attune NxT Software for collecting the data and FlowJo for analyzing the data. Live cells were gated from the FSC/SSC gate for further analysis. Then, the IgM BCR, IgM BCRΔFab cells were gated from GFP and IgM-APC respectively, to show the percentage of the cell populations.

### Protein expression and purification

The stable J558L mouse B cell lines were cultured in Roswell Park Memorial Institute (RPMI) 1640 Medium (Gibco, Catalog number 11875135) with 10% FBS, 100 U/ml Penicillin/Streptomycin, 0.05 mM 2-Mercaptoethanol, 10 mM HEPES (pH 7.5) and then supplemented with 1.2 mM Xanthine (Sigma Aldrich, Catalog number X-4002), 0.11 mM hypoxanthine (Sigma Aldrich, Catalog number H9377) and 1 µg/ml Mycophenolic acid (Sigma Aldrich, Catalog number M3536) for selection of double positive expression. The cells were cultured at 37 °C under 5% CO_2_.

Cell pellets from around 10 L of suspension culture were subjected to 20 cycles of Dounce homogenization in a low-salt buffer containing 20 mM HEPES pH 7.5, 5% glycerol, EDTA-free Protease Inhibitor Cocktail (Sigma Aldrich, Catalog number 04693159001) and Benzonase Nuclease (Sigma Aldrich, Catalog number E8263). The cell lysate was centrifuged at 42,000 rpm for 1 h (45 Ti fixed-angle rotor, Beckman) at 4 °C.

The cell membrane was transferred to solubilization buffer A containing 20 mM HEPES at pH 7.5, 150 mM NaCl, 5% glycerol, 0.5% lauryl maltose neopentyl glycol (LMNG) with 0.05% cholesteryl hemisuccinate (CHS) (Anatrace, Catalog number NG310-CH210), EDTA-free Protease Inhibitor Cocktail, followed by 20 cycles of Dounce homogenization and overnight incubation at 4 °C. Membrane solubilization was followed by centrifugation at 42,000 rpm for 45 min at 4 °C to remove the cell debris and insoluble material. The supernatant was incubated with anti-FLAG M2 affinity gel (Sigma, Catalog number A2220) for 6 h. The beads were washed with 20 column volumes (CV) of buffer B containing 20 mM HEPES at pH 7.5, 150 mM NaCl, 5% glycerol, 0.05% LMNG with 0.005% CHS and eluted by buffer C containing 20 mM HEPES at pH 7.5, 150 mM NaCl, 5% glycerol, 0.005% LMNG with 0.0005% CHS and 100 mg/ml 3xFLAG peptide (ApexBio, Catalog number A6001).

The elution was concentrated to 1 ml by using Amicon Ultra-15 centrifugal filter unit with 100 kDa cutoff (EMD Millipore, Catalog number UFC910024) and loaded to a step gradient of 10%, 20%, 30% and 40% glycerol in buffer D containing 20 mM HEPES at pH 7.5, 150 mM NaCl, 5% glycerol, 0.005% LMNG with 0.0005% CHS, followed by centrifugation for 16 h at 40,000 rpm (MLS-50 swinging-bucket rotor, Beckman). Fractions with target protein of 1 ml were manually collected and concentrated to 200 µl by using Amicon Ultra-0.5 centrifugal filter unit with 100 kDa cutoff (EMD Millipore, Catalog number UFC510024). The protein was fractionated by size-exclusion chromatography (Superose 6 increase 5/150, GE Healthcare) with buffer F (20 mM HEPES at pH 7.5, 150 mM NaCl, 0.004% LMNG and 0.0004% CHS) to remove the glycerol.

The purity and quality of the proteins were characterized at all stages of the purification process using 4%-20% SDS–PAGE (Bio-Rad, Catalog number 4561096) and 3%-12% Bis-Tris gel (ThermoFisher, Catalog number BN1003BOX). The blue native PAGE (BN-PAGE) was run at 150 V for 1 h first, followed by 250 V for another 1 h at 4 °C. The BN-PAGE buffer contained native PAGE running buffer (ThermoFisher, Catalog number BN2001) with the addition of a Coomassie Blue-G250 containing cathode buffer additive (ThermoFisher, Catalog number BN2002). The SDS-PAGE and BN-PAGE were visualized by Coomassie blue staining (Anatrace, Catalog number GEN-QC-STAIN). The expression of all the components including IgM, Igα and Igβ were visualized by western blotting using anti-mouse IgM mu chain (Abcam, Catalog number ab97230), anti-mouse lambda light chain (Novus Biologicals, Catalog number NB7552), anti-FLAG (Sigma Aldrich, Catalog number A8592) and anti-CD79B (Santa Cruz Biotechnology, Catalog number sc-53210) antibodies, respectively.

### Negative staining EM

The peak fractions from gel filtration chromatography were collected and diluted to a final concentration of 0.01 mg/ml for negative staining EM. Copper grids with carbon support film (Electron Microscopy Sciences, Catalog number CF150-CU-UL) were glow discharged for 30 seconds using a Pelco EasyGlow (Ted Pella) instrument. The sample was applied on freshly glow-discharged grids and incubated for 1 min and blotted on filter paper to remove excess buffer. The sample was stained by 6 µL 2% uranyl acetate solution (Electron Microscopy Sciences, Catalog number 22400-2) twice, each for 30 seconds and blotted again on filter paper. Negatively stained samples were imaged on a Joel JEM1400 Transmission Electron Microscope at 120 keV.

### Cryo-EM data collection

The peak fractions from gel filtration chromatography were concentrated to a final concentration of 0.9 mg/ml and crosslinked by 0.4 mM BS(PEG)5 (Thermo Scientific, Catalog number A35396) on ice for 40 min before cryo-EM sample preparation. 3.3 µl of the protein samples were placed onto glow-discharged Quantifoil R1.2/1.3, gold grids with 400 mesh (Electron Microscopy Sciences, Catalog number Q4100AR1.3) before being blotted for 3-3.5 s under 100% humidity at 4 °C and plunged into liquid ethane using a Mark IV Vitrobot (ThermoFisher). Before data collection, all the grids were pre-screened and optimized at Pacific Northwest Center for Cryo-EM at Oregon Health & Science University (PNCC), the University of Massachusetts Cryo-EM Core (UMASS) and Harvard Cryo-EM Center for Structural Biology (HMS) to achieve good ice and particle quality.

Final datasets were collected at HMS using a Titan Krios microscope (ThermoFisher) operating at an acceleration voltage of 300 keV equipped with BioQuantum K3 Imaging Filter (Gatan, slit width 20 eV). Data were collected in two separate sessions. The first session was operated in super resolution mode with 105,000× magnification (0.4125 Å per pixel) and a defocus range between −1.0 and −2.0 µm. For each image stack with 50 frames, the total dose was 55.5 electrons per Å^2^. The second session was operated in super resolution mode with 105,000× magnification (0.415 Å per pixel) and a defocus range between −1.0 and −2.0 µm. For each image stack with 60 frames, the total dose was 60.0 electrons per Å^2^. SerialEM 3.8 was used for fully automated data collection.

### Cryo-EM data processing

The computer support and software for data processing support were provided by SBGrid consortium^34^. For the full-length IgM BCR dataset, raw movies were corrected by gain reference and beam-induced motion by binning two fold with or without dose weighting using the Relion 3.08 implementation of the MotionCor2 algorithm^35^. The motion-corrected micrographs were imported into cryoSPARC^36^ to perform blob picking. For the IgM BCRΔFab dataset, raw movies were corrected by gain reference and beam-induced motion in the same way as for the full-length IgM BCR dataset. The motion-corrected micrographs were imported into cryoSPARC^36^ to perform template picking of 4,496,075 particles using the template generated from the dataset of full-length IgM BCR. Representative 2D classes were then selected as templates for Topaz training^37^. A total number of 3,456,531 particles were picked from a trained Topaz reference resulting in 0.83 per pixel for the individual dataset from two sessions and extracted. Three rounds of 2D classification were performed followed by selecting good particles from good classes. 321,466 particles were selected and merged for further ab initio reconstruction to generate an initial model. These particles from the two sessions were processed similarly for 3D classification into 4 classes. And then the non-uniform refinement with C1 symmetry resulted in 5.9 and 5.4 Å map for two datasets respectively. two classes of 533,389 particles were combined for the final round of 3D classification into 4 classes. 405,695 particles in total were kept for non-uniform refinement. Additional reconstruction using a tight mask led to maps at 3.9 Å resolution. Local refinement on membrane proximal region achieved a 3.6 Å final map. All reported resolutions were estimated based on the gold-standard Fourier shell correlation (FSC) = 0.143 criterion. All the cryo-EM maps were corrected and sharpened by applying a negative B factor using automated procedures in RELION 3.1. Local resolution estimation of all the cryo-EM maps were estimated using Phenix^38^.

### Model fitting and building

We started map interpretation by first building AlphaFold models using the AlphaFold2 implementation in the ColabFold notebooks running on Google Colaboratory^8,9^. Default settings were used with Amber relaxation, and sequences were entered in tandem and separated by a semicolon for prediction of complexes. AlphaFold was run once with each of the 5 trained models, which were checked for consistency. AlphaFold computes pLDDT (predicted local distance difference test, 0-100 with 100 being the best) score to indicate the accuracy of a prediction and we plotted per-residue pLDDT as indication of prediction reliability.

Although several structures of BCR components are available^11,13^, there is no structure for any intact BCR, in particular the full-length structure of the Igα/β heterodimer. To facilitate density interpretation and model building, we first utilized AI-guided protein structure prediction (AlphaFold running on ColabFold notebook)^8,9^ to generate models of BCRΔFab. The prediction gave good per-residue pLDDT scores at the ECD but poorer ones at the TMD. While the overall prediction showed variability among the five ranked models, the predicted ECD and the TMD of the Igα/β heterodimer are highly consistent, and fitted well with the density individually. By contrast, the interaction between mIgM and the Igα/β heterodimer at either the ECD or the TM were predicted poorly. Thus, we manually placed the mIgM ECD and TMD helices separately and built *de novo* the flexible CPs between the ECD and TMD for all chains in COOT^39^. The quality of the map was sufficient for sequence assignments for most of the IgM BCRΔfab complex. Because Cµ2 domain is not visible due to flexibility, the final model of BCRΔFab includes the full-length Igα/β heterodimer and Cµ3-TMD of mIgM. For building the intact BCR, we placed the crystal structures of Fab and CH2 domain^11,12^ into the density. The cytoplasmic tails of Igα and Igβ containing the ITAMs are invisible, supporting their previously recognized dynamic nature^10^, at least in the absence of bound intracellular signalling proteins. Model fitting was conducted using ChimeraX^40^. The BCRΔFab structure was refined by real space refinement followed by model validation in Phenix^38^.

## Acknowledgments

We thank the members of the Wu lab for helpful discussions, R. Walsh, S. Sterling. M. Mayer and S. Rawson at the Harvard Cryo-EM Center for Structural Biology for cryo-EM training and data collection. We also thank J. Myers, V. Rayaprolu and the Pacific Northwest Center for Cryo-EM at Oregon Health & Science University for preliminary dataset collection, under the NIH grant U24GM129547 and accessed through EMSL (grid.436923.9), a DOE Office of Science User Facility sponsored by the Office of Biological and Environmental Research. We thank SBGrid for software and computing support.

## Author contributions

M.R. and H.W. conceptualized the project. F.B., D.S., and J.Y. made the stable cell lines for BCR expression. Z.L. validated BCR expression by flow cytometry under F.W.A.’s supervision. Y.D. resorted and grew the cells, and purified the BCR samples for negative staining and cryo-EM. Y.D. and X.P. performed cryo-EM data acquisition. X.P. and Y.D. performed data processing. Y.D. and X.P. built and refined the model. Y.D. made the figures. H.W., Y.D., and M.R. wrote the manuscript with input from all authors.

## Data and materials availability

All data and materials reported in the main and supplementary data are available upon reasonable requests. The electron density maps will be deposited in the Electron Microscopy Data Bank (EMDB) and the atomic coordinates will be deposited in the Protein Data Bank.

**Extended Data Figure 1.**
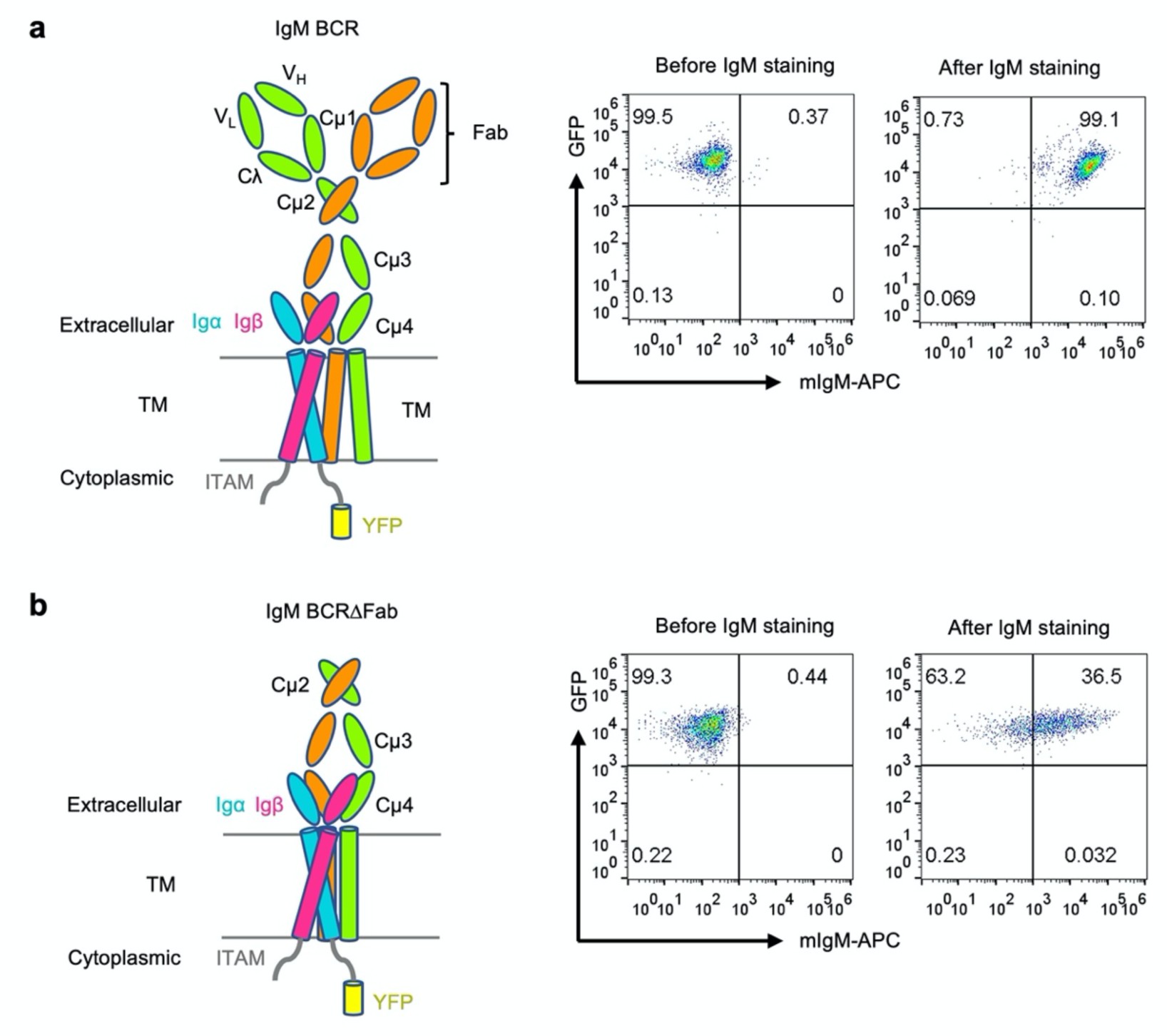
Selection of J558L B cells that co-expressed mIgM and Igα/β for IgM BCR. **(a) and IgM BCRΔFab (b)**. Left: domain organization. Right: flow cytometry and cell sorting by surface staining of IgM and using GFP channel for the YFP in Igα.

**Extended Data Figure 2.**
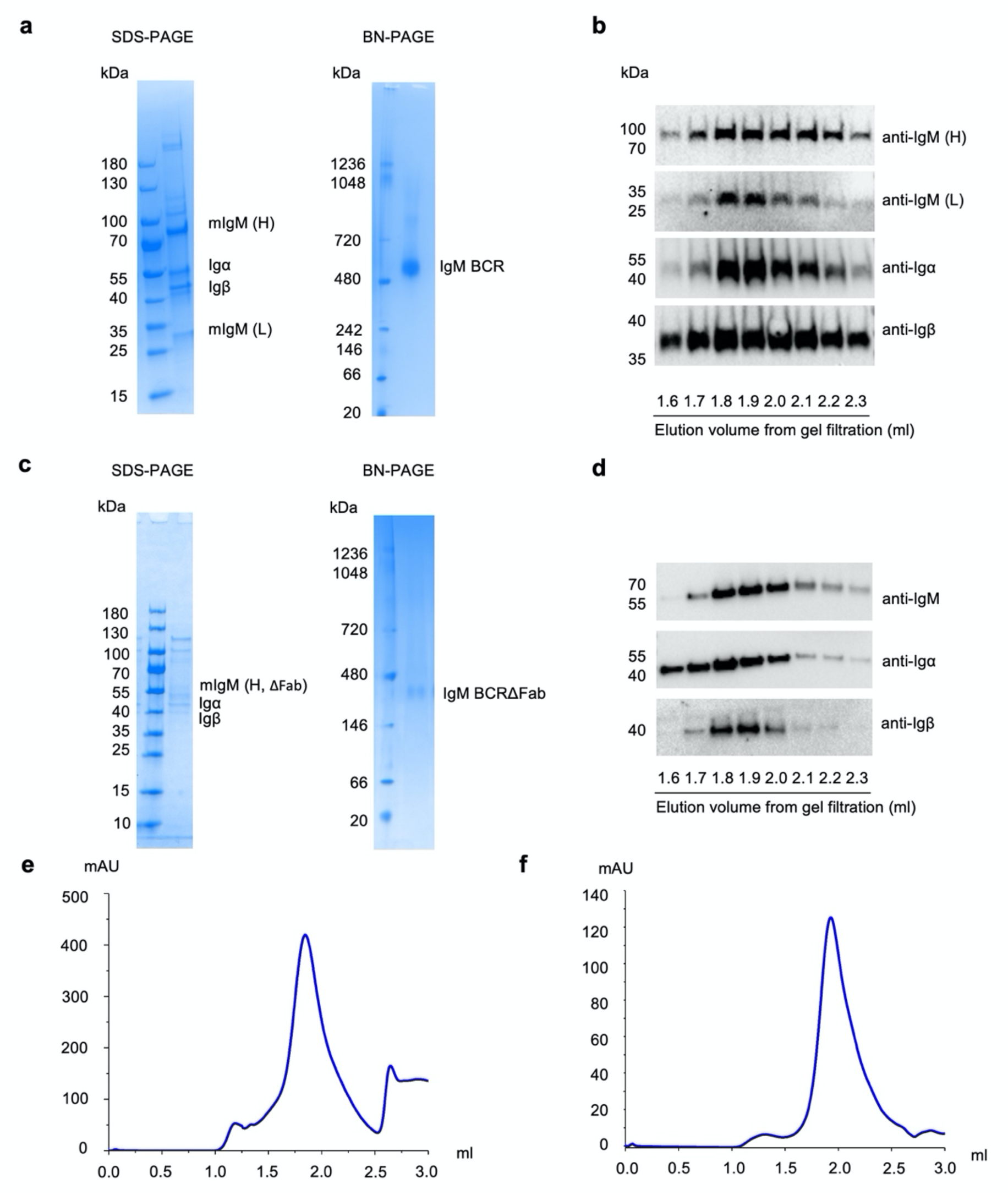
Protein purification of IgM BCR and IgM BCRΔFab. **a-d**, Purification of IgM BCR (a, b) and IgM BCRΔFab (c, d) by SDS-PAGE, Blue-native (BN) PAGE and western blot using antibodies against the individual subunits. IgM (H): heavy chain; IgM (L): light chain. **e-f**, Gel filtration profile of IgM BCR (e) and IgM BCRΔfab (f).

**Extended Data Figure 3.**
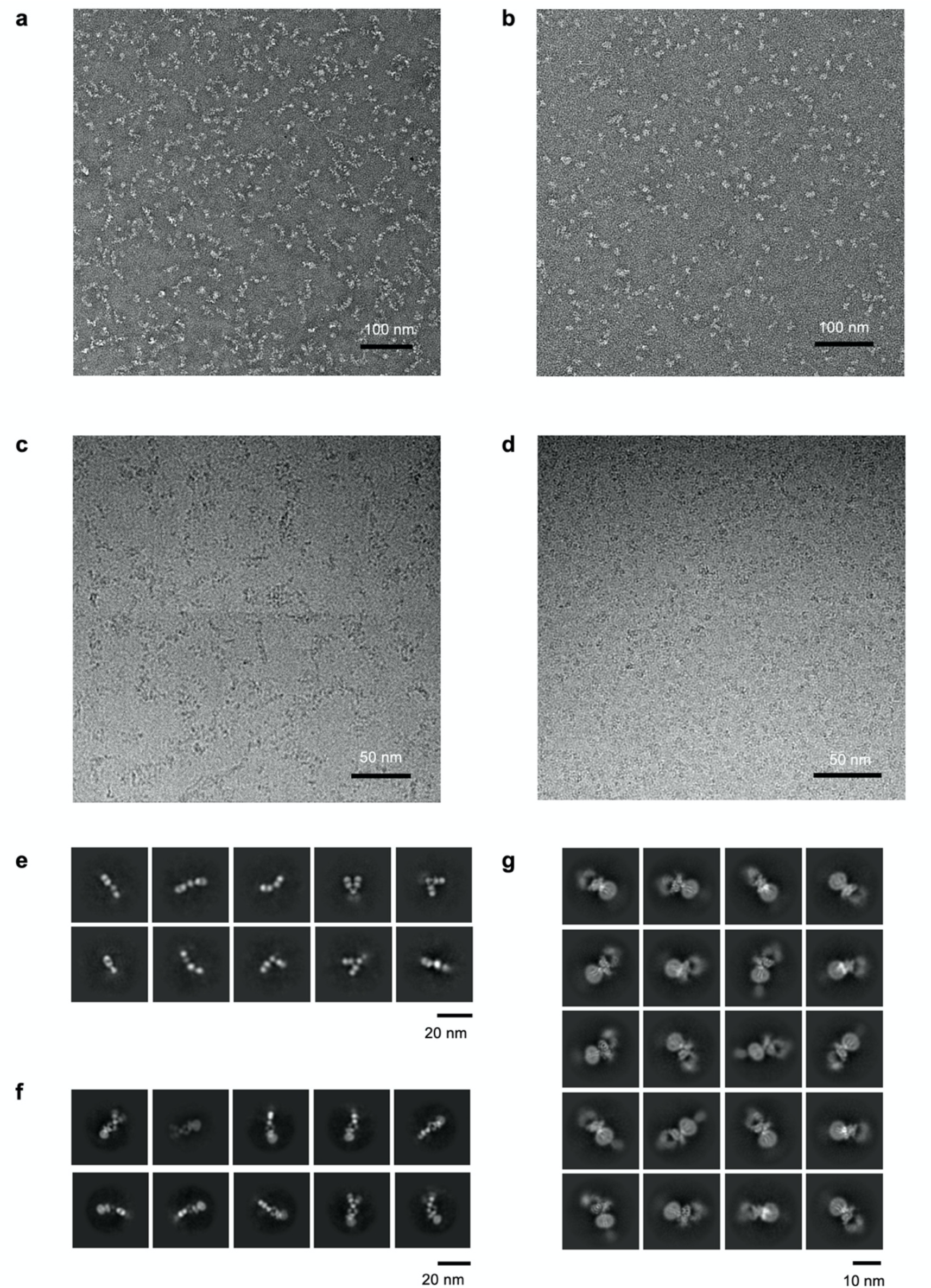
Negative staining raw images, cryo-EM raw images and 2D classifications. **a-b**, Representative negative staining EM images of IgM BCR (a) and IgM BCRΔFab (b). **c-d**, Representative cryo-EM images of IgM BCR (c) and IgM BCRΔFab (d). **e**, Representative 2D classes of IgM BCR specifically at its Fab region, showing a large-scale conformational change. **f-g**, Representative 2D classes of the overall IgM BCR and IgM BCRΔFab.

**Extended Data Figure 4.**
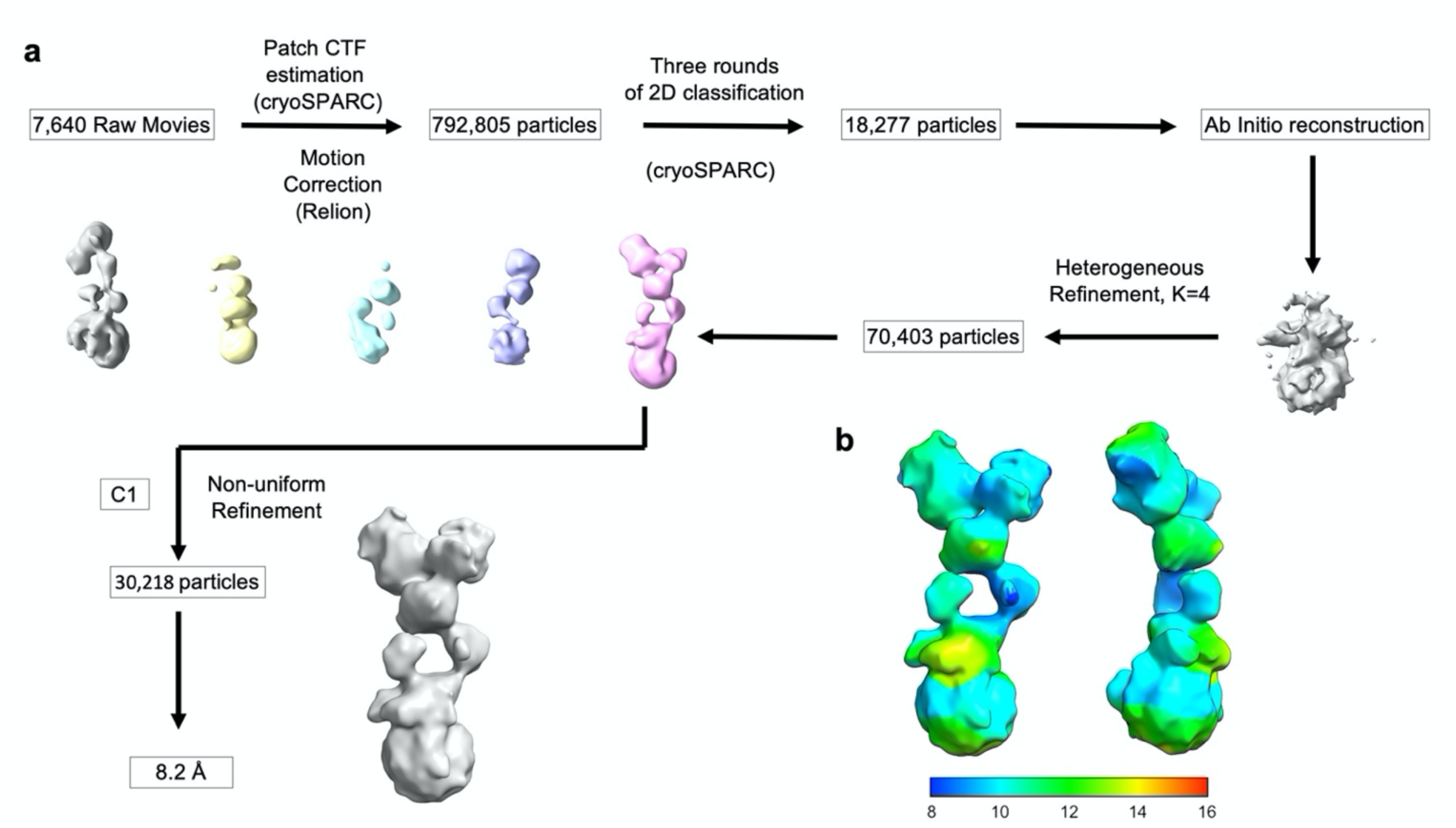
Cryo-EM structure determination of IgM BCR. **a**, Data processing flow chart. **b**, Local resolution distribution.

**Extended Data Figure 5.**
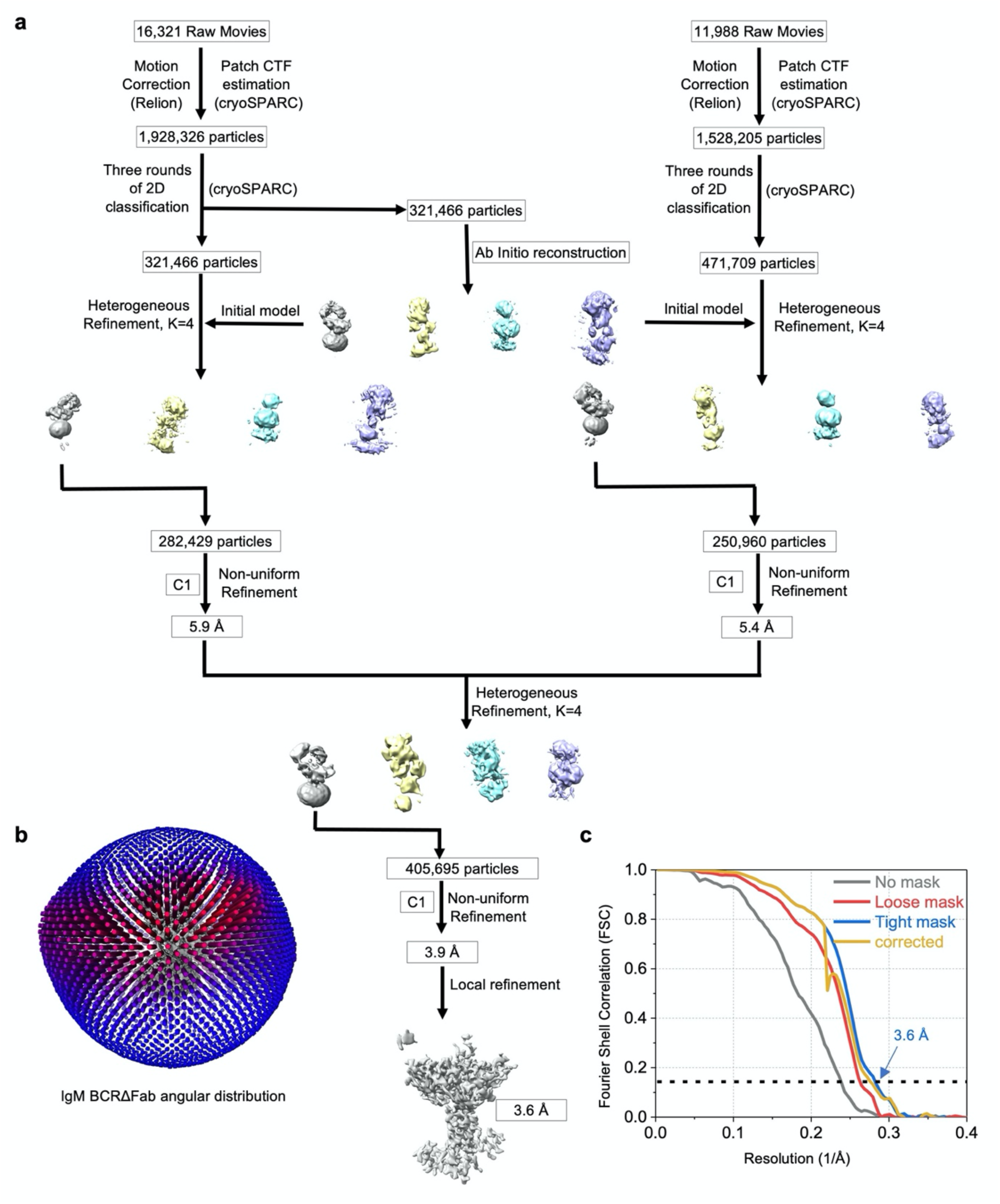
Cryo-EM data processing for IgM BCRΔFab. **a**, cryo-EM data processing flow chart. **b**, Angular distribution of the particles used for the final reconstruction. **c**, Fourier shell correlation plots.

**Extended Data Figure 6.**
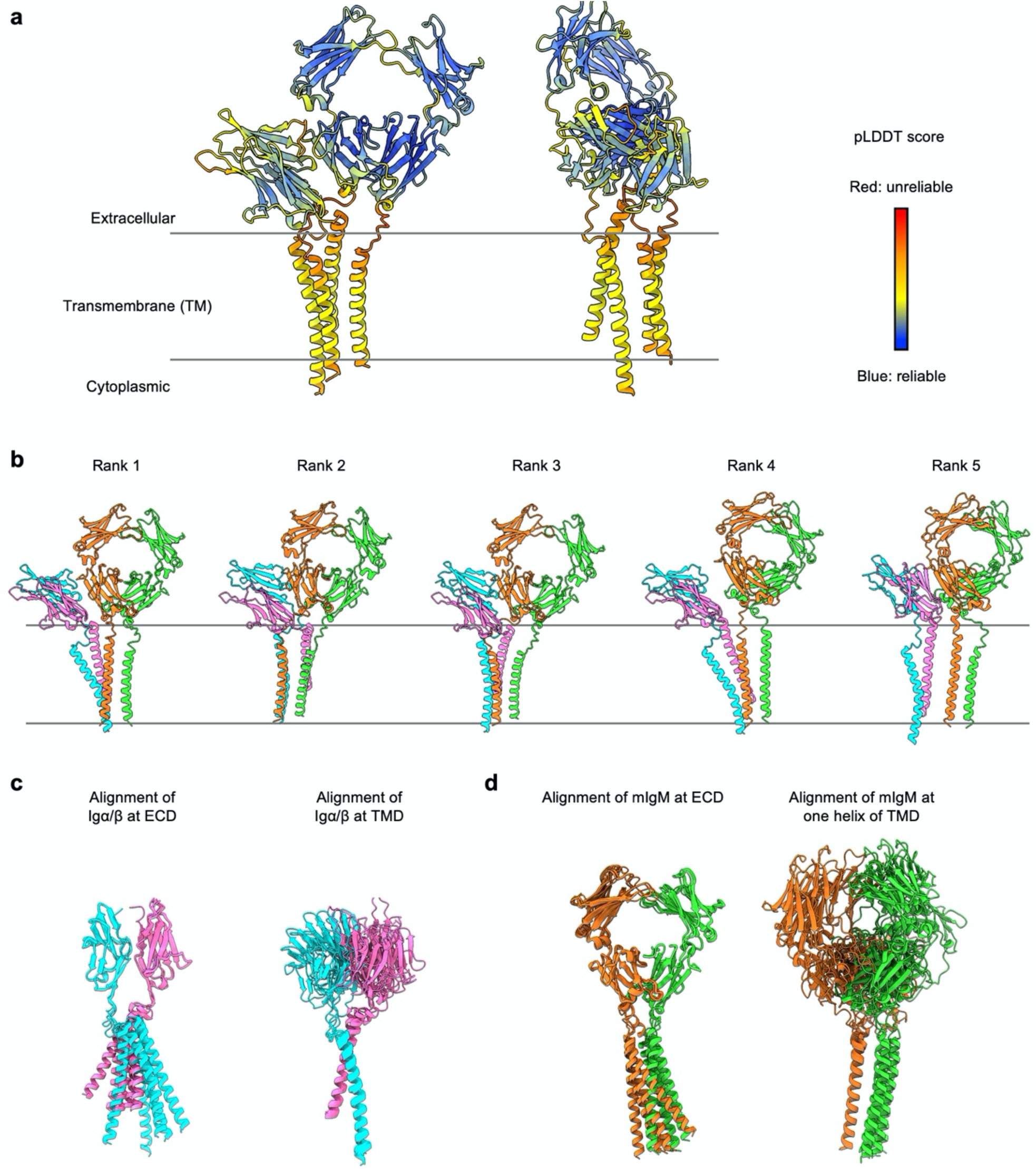
AlphaFold predicted model of IgM BCR. **a**, The top ranked model of IgM BCR (without Fab and Cµ2) from AlphaFold, coloured by per-residue pLDDT score. **b**, Five predicted models of IgM BCR are shown side by side. **c**, Alignment of the five models of the Igα/β heterodimer at ECD (left) and TMD (right), showing consistent prediction of the ECD interaction (left) and the TMD interaction (right) separately. The CPs of the Igα/β heterodimer were not predicted correctly. **d**, Alignment of the five models of mIgM at ECD (left) and TMD (right), showing consistent prediction of the ECD interaction (left) but lack of consistency in the prediction of the TMD interaction (right). The CPs of the mIgM dimer were not predicted correctly.

**Extended Data Figure 7.**
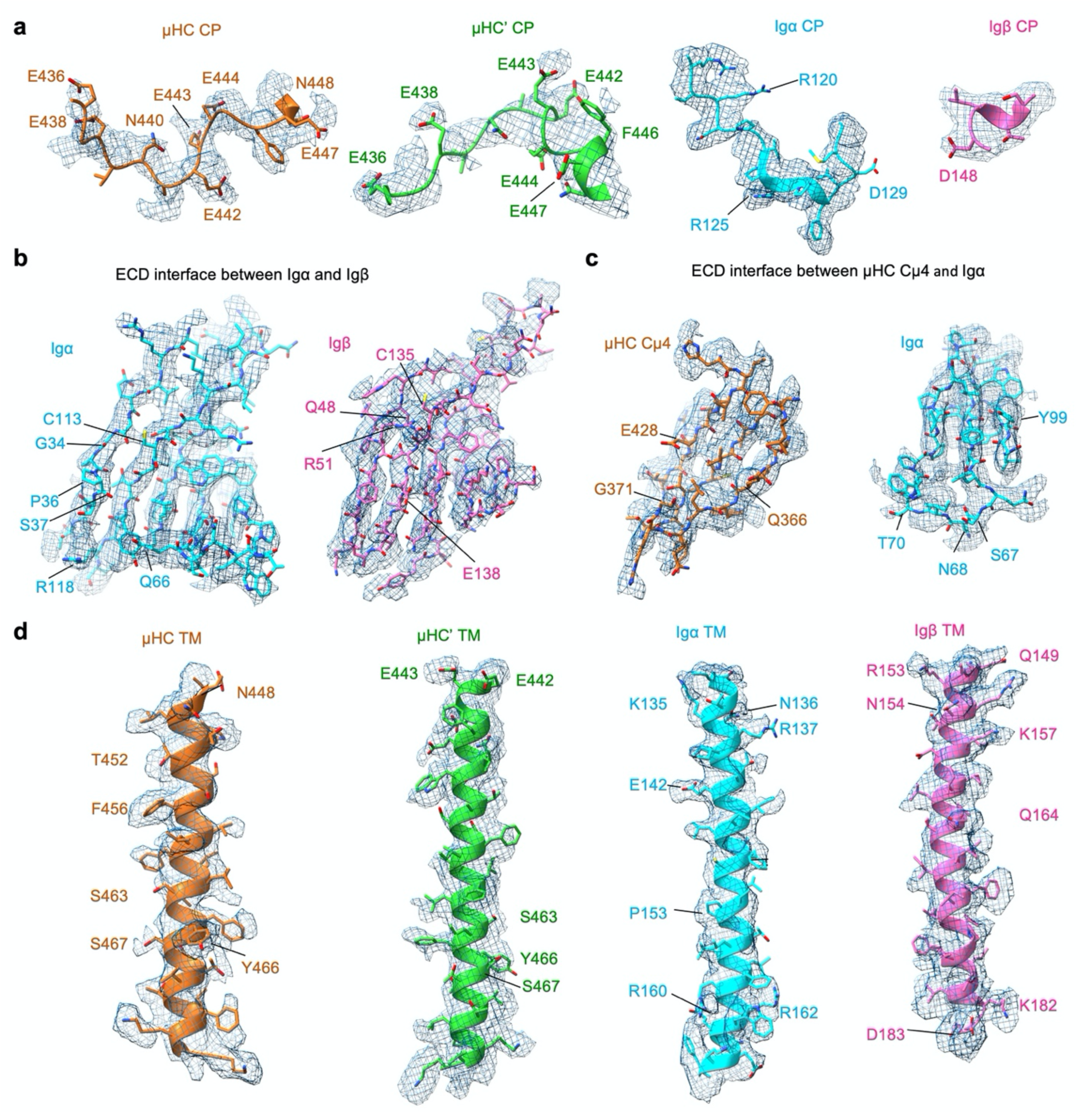
Quality of model fitting in the IgM BCRΔFab cryo-EM map (contour level: 8.5 σ). **a**, Four connecting peptide regions from µHC, µHC’, Igα and Igβ superimposed with the cryo-EM map. Acidic and basic residues are shown and labeled. **b**, Ig regions of Igα and Igβ for the Igα/β interaction superimposed with the cryo-EM map. **c**, Interface between µHC Cµ4 and Igα superimposed with the cryo-EM map. **d**, TMD helices of µHC, µHC’, Igα and Igβ superimposed with the cryo-EM map. Key residues mediating the TMD interaction are shown.

**Extended Data Figure 8.**
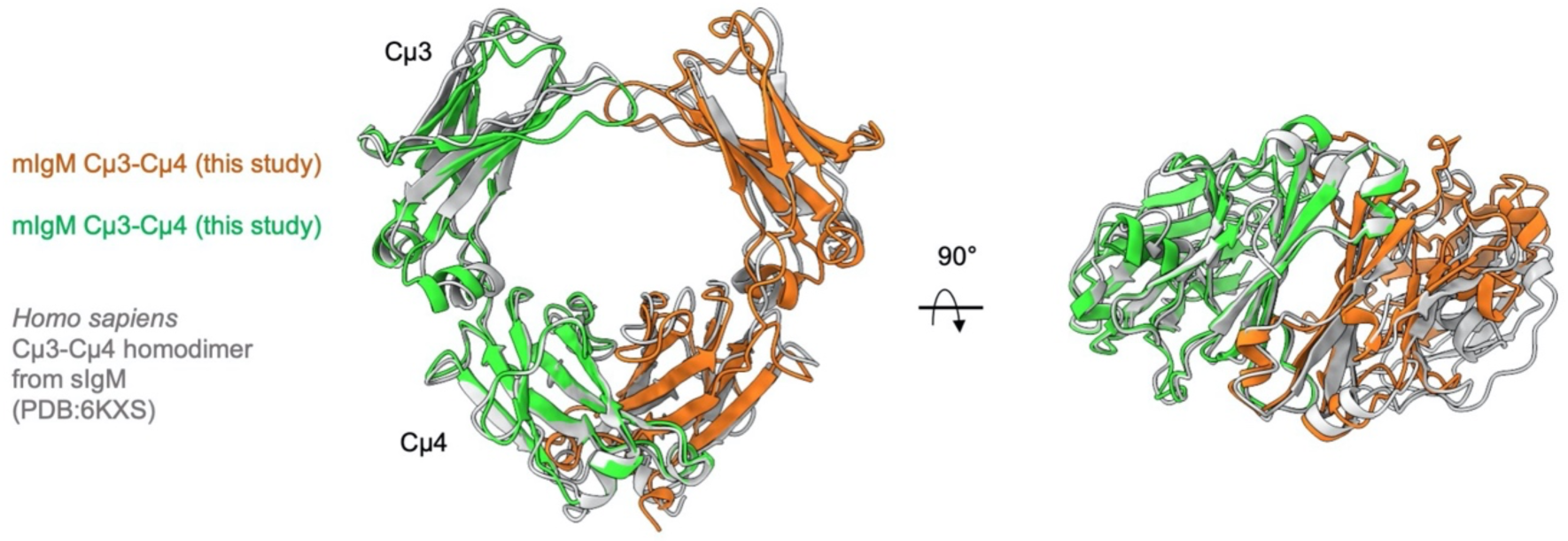
Alignment of the Cµ3-Cµ4 homodimer of IgM BCR with that from the crystal structure of secreted IgM or sIgM (PDB: 6KXS).

**Extended Data Figure 9.**
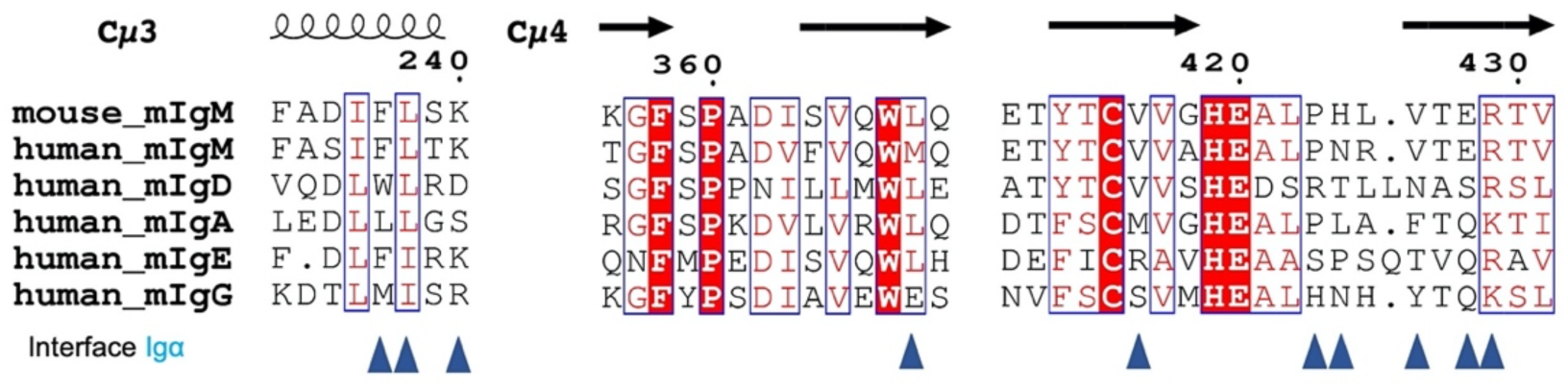
Residues in mIgM ECD for Igα interaction are not conserved in other mIg isotypes. Sequence alignment of part of the Cµ3-Cµ4 domains with other mIg isoforms. Residues at the interface with Igα are shown by triangle symbols.

**Extended Data Figure 10.**
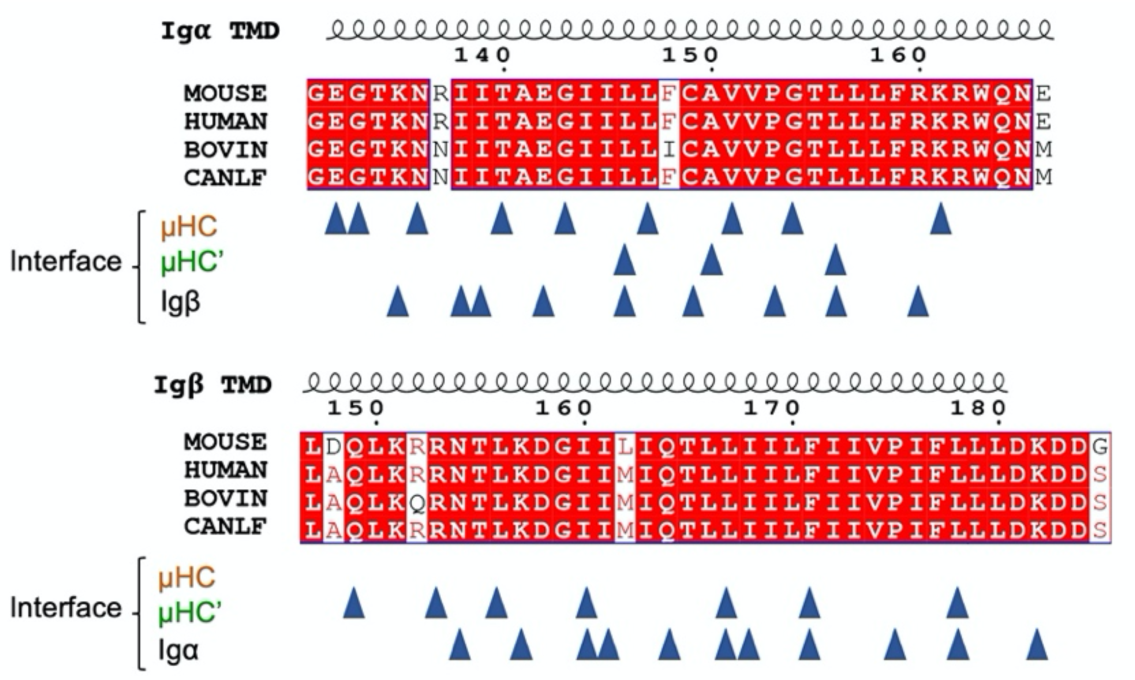
Sequence alignment of Igα and Igβ TMD among the indicated species, with the interactions shown by triangles. Residues at the interface of Igα and Igβ with µHC, µHC’, and Igα or Igβ are indicated. µHC TMD does not interact with Igβ TMD. CANLF: dog.

**Extended Data Figure 11.**
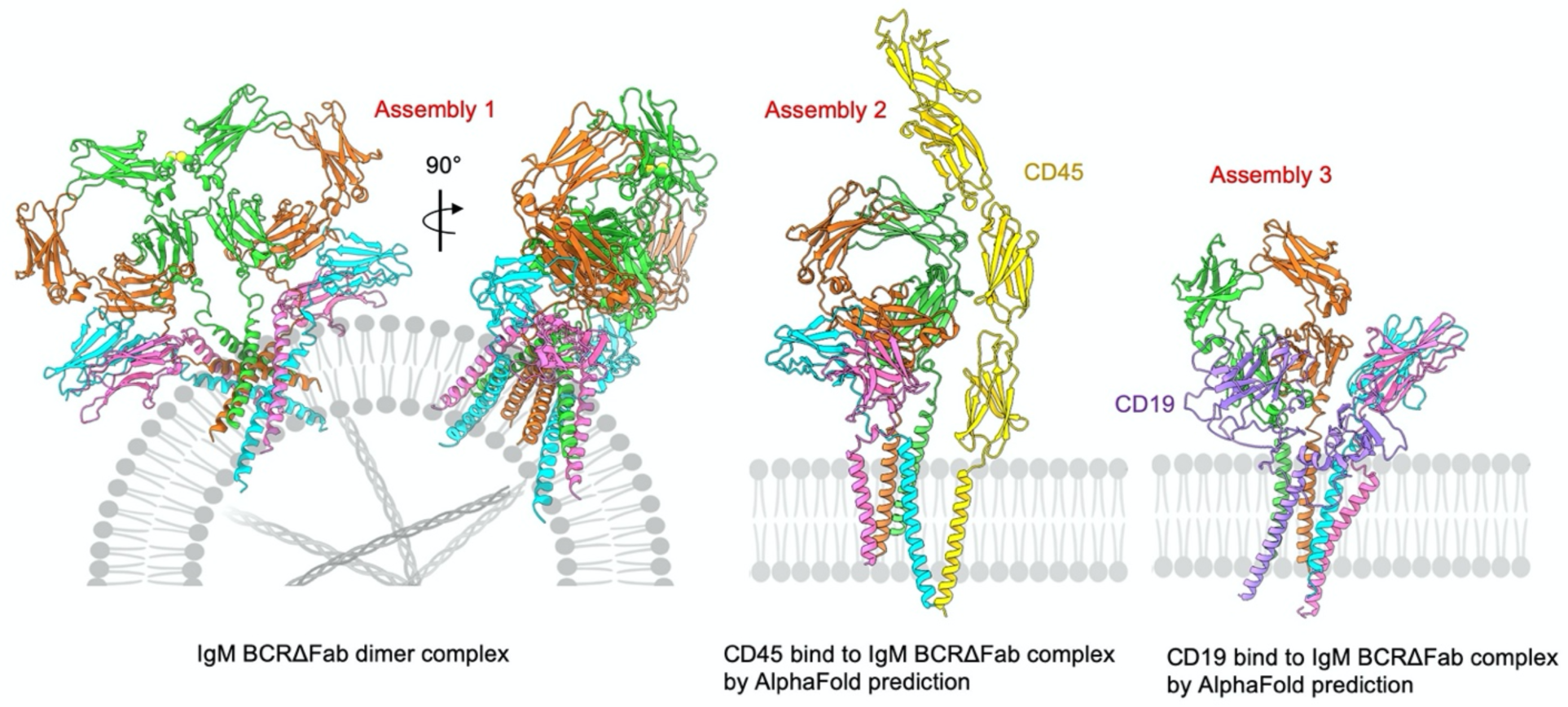
Proposed models for dynamic assembly of IgM BCR with other signalling components. Assembly 1 is an IgM BCR dimer generated by docking a second BCR using pentameric sIgM (PDB:6KXS) as a reference. Assembly 2 and 3 were taken from models predicted by AlphaFold.

**Table S1.**
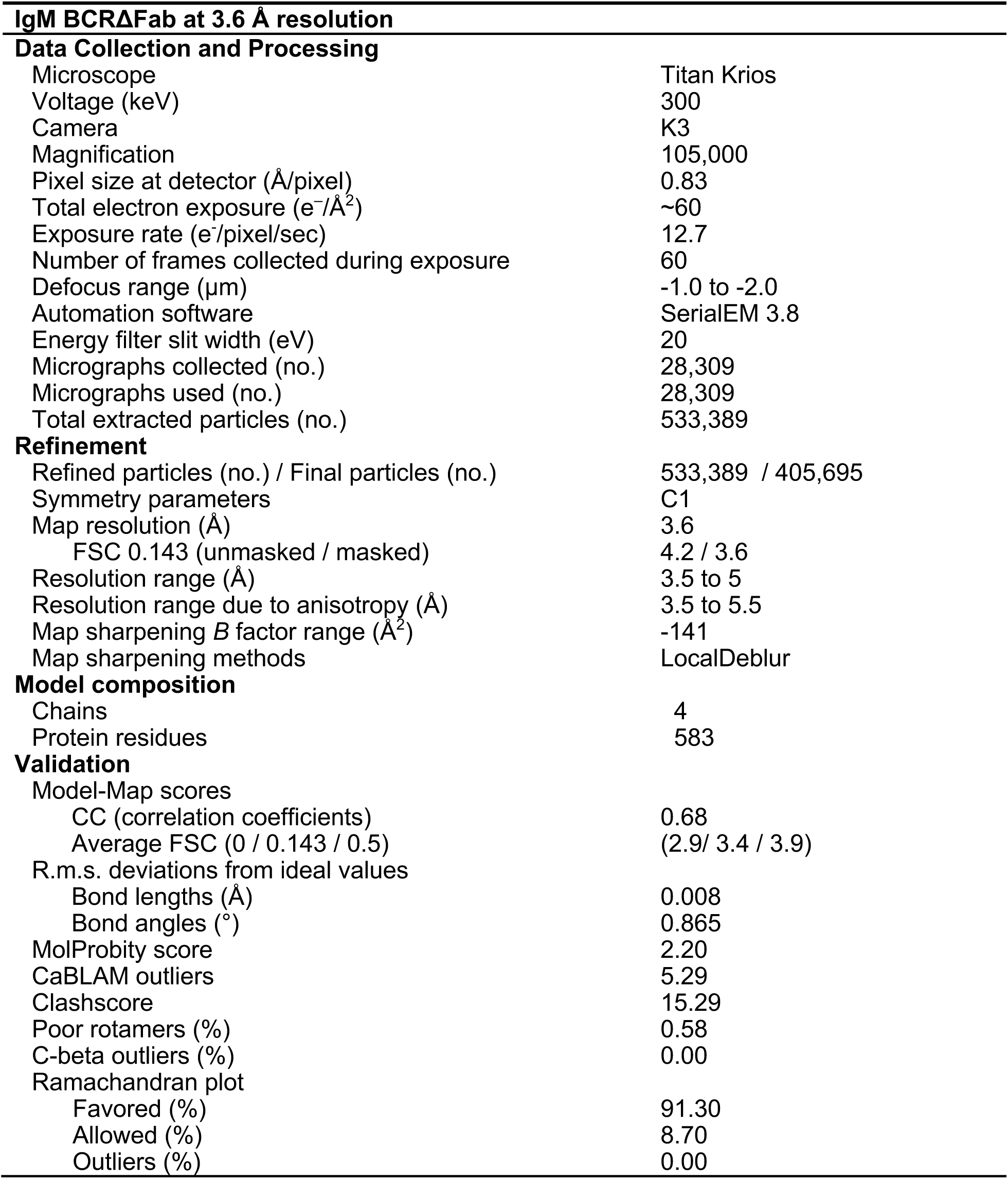
Data Collection, data processing and validation statistics

## References

1. Bengtén, E. et al. Immunoglobulin Isotypes: Structure, Function, and Genetics. in Origin and Evolution of the Vertebrate Immune System (eds. Du Pasquier, L. & Litman, G. W.) vol. 248 189–219 (Springer Berlin Heidelberg, 2000).

2. Stavnezer, J. & Schrader, C. E. Ig heavy chain class switch recombination: mechanism and regulation. J. Immunol. Baltim. Md 1950 193, 5370–5378 (2014).

3. Schroeder, H. W. & Cavacini, L. Structure and Function of Immunoglobulins. J. Allergy Clin. Immunol. 125, S41–S52 (2010).

4. Roux, K. H., Strelets, L., Brekke, O. H., Sandlie, I. & Michaelsen, T. E. Comparisons of the Ability of Human IgG3 Hinge Mutants, IgM, IgE, and IgA2, to Form Small Immune Complexes: A Role for Flexibility and Geometry. J. Immunol. 161, 4083–4090 (1998).

5. Adlersberg, J. B. The immunoglobulin hinge (Interdomain) region. Ric. Clin. Lab. 6, 191 (1976).

6. Sandin, S., Öfverstedt, L.-G., Wikström, A.-C., Wrange, Ö. & Skoglund, U. Structure and Flexibility of Individual Immunoglobulin G Molecules in Solution. Structure 12, 409–415 (2004).

7. Wurzburg, B. A., Garman, S. C. & Jardetzky, T. S. Structure of the Human IgE-Fc C⑀3-C⑀4 Reveals Conformational Flexibility in the Antibody Effector Domains. 11.

8. Mirdita, M. et al. ColabFold - Making protein folding accessible to all. 8.

9. Jumper, J. et al. Highly accurate protein structure prediction with AlphaFold. Nature 596, 583–589 (2021).

10. Dong, D. et al. Structural basis of assembly of the human T cell receptor–CD3 complex. Nature 573, 546–552 (2019).

11. Müller, R. et al. High-resolution structures of the IgM Fc domains reveal principles of its hexamer formation. Proc. Natl. Acad. Sci. 110, 10183–10188 (2013).

12. Ban, N. et al. Structure of an anti-idiotypic Fab against feline peritonitis virus-neutralizing antibody and a comparison with the complexed Fab. FASEB J. 9, 107–114 (1995).

13. Radaev, S. et al. Structural and Functional Studies of Igαβ and Its Assembly with the B Cell Antigen Receptor. Structure 18, 934–943 (2010).

14. Krissinel, E. & Henrick, K. Inference of Macromolecular Assemblies from Crystalline State. J. Mol. Biol. 372, 774–797 (2007).

15. Li, Y. et al. Structural insights into immunoglobulin M. Science 367, 1014–1017 (2020).

16. Hiramoto, E. et al. The IgM pentamer is an asymmetric pentagon with an open groove that binds the AIM protein. Sci. Adv. 4, eaau1199 (2018).

17. Vuillier, F. et al. Lower levels of surface B-cell-receptor expression in chronic lymphocytic leukemia are associated with glycosylation and folding defects of the μ and CD79a chains. Blood 105, 2933–2940 (2005).

18. Campbell, K. S., Hager, E. J. & Cambier, J. C. Alpha-chains of IgM and IgD antigen receptor complexes are differentially N-glycosylated MB-1-related molecules. 7.

19. Schamel, W. W. A. & Reth, M. Stability of the B cell antigen receptor complex. Mol. Immunol. 37, 253–259 (2000).

20. Schwans, J. P. et al. Use of anion–aromatic interactions to position the general base in the ketosteroid isomerase active site. Proc. Natl. Acad. Sci. 110, 11308–11313 (2013).

21. Kwong, P. D. et al. Structure of an HIV gp120 envelope glycoprotein in complex with the CD4 receptor and a neutralizing human antibody. Nature 393, 648–659 (1998).

22. Schamel, W. W. A. & Reth, M. Monomeric and Oligomeric Complexes of the B Cell Antigen Receptor. Immunity 13, 5–14 (2000).

23. Tolar, P., Sohn, H. W. & Pierce, S. K. The initiation of antigen-induced B cell antigen receptor signaling viewed in living cells by fluorescence resonance energy transfer. Nat. Immunol. 6, 1168–1176 (2005).

24. Gottwick, C. et al. A symmetric geometry of transmembrane domains inside the B cell antigen receptor complex. Proc. Natl. Acad. Sci. 116, 13468–13473 (2019).

25. Ghosh, M. R., Kim, B. S. & Tucker, P. W. Differential Structure-Function Requirements of the Transmembranal Domain of the B Cell Antigen Receptor By V. S. Parikh,* G. A. Bishop,∼ K.-J. Liu,$ B. T. Do,. 7 (1992).

26. Shaw, C., Mitchell, N., Weaver, Y. K. & Abbas, A. K. Mutations of lmmunoglobulin Transmembrane and Cytoplasmic Domains: Effects on Intracellular Signaling and Antigen Presentation. 12.

27. Friess, M. D., Pluhackova, K. & Böckmann, R. A. Structural Model of the mIgM B-Cell Receptor Transmembrane Domain From Self-Association Molecular Dynamics Simulations. Front. Immunol. 9, 2947 (2018).

28. Yang, J. & Reth, M. Oligomeric organization of the B-cell antigen receptor on resting cells. Nature 467, 465–469 (2010).

29. Gold, M. R. & Reth, M. G. Antigen Receptor Function in the Context of the Nanoscale Organization of the B Cell Membrane. Annu. Rev. Immunol. 37, 97–123 (2019).

30. DeFord-Watts, L. M. et al. The Cytoplasmic Tail of the T Cell Receptor CD3 ε Subunit Contains a Phospholipid-Binding Motif that Regulates T Cell Functions. J. Immunol. 183, 1055–1064 (2009).

31. Zimmerberg, J. & McLaughlin, S. Membrane Curvature: How BAR Domains Bend Bilayers. Curr. Biol. 14, R250–R252 (2004).

32. Mim, C. & Unger, V. M. Membrane curvature and its generation by BAR proteins. Trends Biochem. Sci. 37, 526–533 (2012).

33. Saltukoglu, D. et al. Plasma membrane topography governs the three-dimensional dynamic localization of IgM B cell receptor clusters. http://biorxiv.org/lookup/doi/10.1101/2022.04.29.489661 (2022) xdoi:10.1101/2022.04.29.489661.

34. Morin, A. et al. Collaboration gets the most out of software. eLife 2, e01456 (2013).

35. Zheng, S. Q. et al. MotionCor2 - anisotropic correction of beam-induced motion for improved cryo-electron microscopy. Nat. Methods 14, 331–332 (2017).

36. Punjani, A., Rubinstein, J. L., Fleet, D. J. & Brubaker, M. A. cryoSPARC: algorithms for rapid unsupervised cryo-EM structure determination. Nat. Methods 14, 290–296 (2017).

37. Bepler, T. et al. Positive-unlabeled convolutional neural networks for particle picking in cryoelectron micrographs. Nat. Methods 16, 1153–1160 (2019).

38. Adams, P. D. et al. PHENIX: a comprehensive Python-based system for macromolecular structure solution. Acta Crystallogr. D Biol. Crystallogr. 66, 213–221 (2010).

39. Emsley, P. & Cowtan, K. Coot : model-building tools for molecular graphics. Acta Crystallogr. D Biol. Crystallogr. 60, 2126–2132 (2004).

40. Goddard, T. D. et al. UCSF ChimeraX: Meeting modern challenges in visualization and analysis. Protein Sci. Publ. Protein Soc. 27, 14–25 (2018).

